# Effort Foraging Task reveals positive correlation between individual differences in the cost of cognitive and physical effort in humans

**DOI:** 10.1101/2022.11.21.517394

**Authors:** Laura A. Bustamante, Temitope Oshinowo, Jeremy R. Lee, Elizabeth Tong, Allison R. Burton, Amitai Shenhav, Jonathan D. Cohen, Nathaniel D. Daw

## Abstract

Effort-based decisions, in which people weigh potential future rewards against effort costs required to achieve those rewards, have largely been studied separately for cognitive or physical effort, yet most real-world actions incur both effort costs. What is the relationship between cognitive and physical effort costs? Here we attempt to formalize the mechanisms underlying effort-based decisions and address methodological challenges to isolate and measure these mechanisms. Patch foraging is an ecologically valid reward rate maximization problem with well-developed theoretical tools to understand choices. We developed the Effort Foraging Task, which embedded cognitive or physical effort into a patch foraging sequential decision task, to isolate and quantify the cost of both cognitive and physical effort using a computational model. Participants chose between harvesting a depleting patch, or traveling to a new patch that was costly in time and effort. Participants’ exit thresholds (reflecting the reward they expected to receive by harvesting when they chose to travel to a new patch) were sensitive to cognitive and physical effort demands, allowing us to quantify the perceived effort cost in monetary terms. Individual differences in cognitive and physical effort costs were positively correlated, suggesting that these are perceived and processed in common terms across different domains. We found patterns of correlation of both cognitive and physical effort costs with self-reported anxiety, cognitive function, behavioral activation, and self-efficacy. This suggests the task captures decision mechanisms associated with real-world motivation and can be used to study individual variation in effort-based decisions across domains of cost.

## 1 Introduction

People make *effort-based decisions* every day, weighing the potential rewards associated with an action against the effort it requires. These decisions can involve cognitive effort, physical effort, or both. Economic utility theory has been productively applied in effort-based decision making research: people seek to maximize reward while minimizing effort, which can be accomplished by computing an ‘expected value’ of effort (Walton et al., 2007; Rigoux & Guigon, 2012; Shenhav et al., 2013; Shenhav et al., 2017; Salamone et al., 2018). In these theories effort is described as costly, reducing the value of rewards. Evidence from cognitive psychology and neuroscience shows that people consistently factor effort into their decisions but that individuals approach tradeoffs between rewards and cognitive (Kool et al., 2010; Westbrook et al., 2013) and physical (Treadway et al., 2009; Treadway, Buckholtz, et al., 2012) effort differently. Effort related behaviors range from effort seeking (e.g., running a marathon, writing a book) to effort avoiding (e.g., sedentary behavior, academic procrastination). There is much to learn about the trait and state factors that lead to effort seeking versus avoiding. Advancing knowledge of effort-based decision making is important because of evidence demonstrating proxies of cognitive and physical effort costs are related to psychiatric symptoms, including apathy and anhedonia in depression (Cĺery-Melin et al., 2011; Treadway, Bossaller, et al., 2012; Yang et al., 2014; Zald & Treadway, 2017; Grahek et al., 2018; Marchetti et al., 2018; Patzelt et al., 2019; Tran et al., 2021; Westbrook et al., 2022) and negative symptoms of schizophrenia (Barch et al., 2014; Wolf et al., 2014; Culbreth et al., 2016). What are the shared versus distinct contributions of cognitive and physical effort costs to these symptoms?

Reinforcement learning literature has been concerned with the extent to which there is a common representation of ‘value’ that integrates different kinds of rewards (e.g., food, money Chib et al., 2009; Levy & Glimcher, 2011, 2012). To what extent is there also a common representation of ‘cost’ that integrates different domains of effort and other costs associated with an activity (i.e. cognitive and physical effort, and time costs, Schmidt et al., 2012; Vinckier et al., 2022)? Relatedly, how are effort costs common or different across the wide range of tasks that can be considered “physical” or “cognitive”? Do “cognitive” and “physical” effort constitute separate domains, or is there a different organizational principle?

Human and animal research suggests that cognitive and physical effort-based decisions are controlled by shared neural populations (Schmidt et al., 2012; Chong et al., 2017; Borderies et al., 2020; Bornert & Bouret, 2021; Lopez-Gamundi et al., 2021). Individual differences provide a window into the relationship between different domains of cost. Do individuals who avoid cognitive effort more also avoid physical effort more? Research is limited about the relationship between individual differences in cognitive and physical effort costs. Lopez-Gamundi & Wardle (2018) found a positive relationship (r=0.43) between the percentage of ‘hard task choices’ in the cognitive (task-switching task) and physical (rapid key-pressing task) versions of the Effort Expenditure for Rewards Task (this finding was replicated in Tran et al., 2020, r=0.35).

We offer a new approach to these questions by making use of a novel variant of a decision task from the foraging literature, which offers a distinct approach to studying how organisms integrate rewards and costs. The study of foraging using serial decision problems originated in ethology and is noteworthy for strong theoretical foundations and ecological validity (Stephens & Krebs, 1986). More recently, it has been increasingly adopted in psychology and neuroscience (Todd et al., 2012; Hills et al., 2015; Hayden & Niv, 2021). In classic ecological contexts, foraging tasks study how organisms optimize some fitness objective (e.g. maximizing reward rate), while balancing rewards (e.g., food) and costs (especially time). Foraging-style tasks have proven to be valuable in understanding decision making in formally rigorous terms, and relating it to underlying neural mechanisms, across a variety of species, including rodents (Carter & Redish, 2016; Kane et al., 2021), non-human primates (Hayden et al., 2011; Hayden, 2018) and humans (Hills et al., 2008; Hills et al., 2012; Kolling et al., 2012; Constantino et al., 2017; Lenow et al., 2017).

### The Effort Foraging Task

In the present study we developed a novel ‘Effort Foraging Task’ designed to leverage the strengths of the patch foraging paradigm (i.e., ecologically valid serial decisions, well-developed formal frameworks, and common neural and cognitive substrates across a range of settings, Stephens & Krebs, 1986; Hills et al., 2008; Todd et al., 2012; Blanchard & Hayden, 2015; Carter et al., 2015; Hills et al., 2015; Carter & Redish, 2016). The task had two variants — cognitive and physical — which we used to fit a computational model to individual participants’ behavior to estimate the costs associated with each form of effort. Within this model we evaluated the correlation between the estimated individual differences in cognitive and physical effort costs.

The Effort Foraging Task adapts a version of the computerized patch-foraging task developed by Constantino & Daw (2015), adding travel costs in the form of cognitively and physically effortful tasks. In a standard patch foraging task, the forager visits a ‘patch’ which can be harvested to yield rewards (here, a simulated orchard with apple trees). Within a given patch, the marginal return (apples) associated with each successive harvest decreases over time. At any point the forager can travel to a new patch, which has replenished rewards, but it takes time to travel there. Deciding when to leave a depleting patch in a foraging environment involves tradeoffs between harvesting rewards available from the current patch, and the time spent traveling to a different (but richer) one. For this reason, the level at which the forager decides to exit the current patch (i.e., their ‘exit threshold’ or the number of apples they have last received before quitting) reflects the reward they are willing to forgo by leaving that patch and spending the time to travel to another. In these respects, the exit threshold reveals the point of equivalence in the tradeoff between the cost of harvesting with diminishing rewards and the time cost of traveling to a new patch.

These considerations are formalized in the Marginal Value Theorem (Charnov, 1976), which asserts that a simple threshold policy maximizes reward rate. The forager simply needs to maintain an estimate of the average reward rate in the environment and exit a patch when the instantaneous reward rate falls below the long run average.

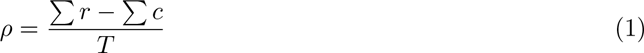

According to equation 1, reward rate is maximized when the exit threshold (*ρ*, the reward level at which the forager exits), is equal to the long run average reward rate, which includes the sum of all rewards (Σ*r*) minus sum of (non-time) costs incurred in the environment (Σ*c*, e.g., energy spent extracting rewards or traveling to the next patch), divided by the total number of harvest periods (the time cost normalized by the harvest time, *T* = total time / harvest time).

Constantino & Daw (2015) found that human participants playing a virtual foraging game used a threshold exit strategy consistent with the Marginal Value Theorem, which explained behavior better than other reinforcement learning models (e.g., temporal difference learning). Furthermore, they found that exit thresholds (the reward expected to receive by harvesting when participants chose to travel to a new patch) shifted reliably and in predicted ways when the environment changed (e.g., when travel time and/or reward depletion was experimentally manipulated). For example, when the travel time between patches was increased, participants’ exit thresholds decreased, reflecting the increased opportunity cost of travel time and an overall decrease in average reward rate.

For the Effort Foraging Task, rather than manipulating travel time, we manipulated travel costs by varying the *effort* – cognitive or physical – required to travel between patches and compared exit thresholds in high versus low effort conditions. In line with MVT, which predicts an increase in travel costs to result in a lower the exit threshold, we predicted that contexts with higher effort costs would, in general, decrease participants’ estimates of average reward rate, leading the exit threshold to be lower; that is, a greater willingness to accept diminishing rewards to avoid effortful travel. Conversely, some participants might instead exhibit effort-seeking, i.e., treat the low effort task as more costly, exiting at a lower threshold in low compared to high effort conditions. Accordingly, we used the difference in exit threshold between high and low effort conditions to infer the perceived costs of travel.

More specifically, we used participants’ decision thresholds to create a computational model based on the Marginal Value Theorem to quantify the added cost of high compared to low effort conditions in this task. Using this model, we found that most participants avoided the high effort tasks, treating them as costly, while some participants sought high-effort tasks, treating them as valued. We to directly fit the correlation between individual differences in cognitive and physical effort costs in the same currency (money) and found a moderate positive correlation.

To assess the external validity of these cost measures, we used a data driven approach — canonical correlation analysis — to look for relationships between task behavior and self-report surveys of motivation and affect (including measures of apathy, anhedonia, depression, and others). These results demonstrated the value of measuring multiple effort domains within-participant, as we found that although individual differences in cognitive and physical effort cost were positively correlated, they had differential relationships to self-reported real-world behavior.

In developing the task, we conducted four experiments. Experiment 1 (N=678) is the focus of our results as it has the largest sample size and is the most efficient version to administer (16 minutes of main task time per effort type). Experiments 2-4 were conducted largely to validate results and rule out confounds. Specifically, Experiment 2 (N=116) serves to demonstrate the generalizability of the method to multiple cognitive effort tasks exercising multiple cognitive processes (interference control, in Experiment 1, working memory in Experiment 2); Experiment 3 (N=43) verifies that standard patch-foraging manipulations are effective in the novel effort context (thresholds sensitive to patch richness); and Experiment 4 (N=71) addresses the possibility that effort avoidance might be confounded by differences in subjective travel duration perception across effort levels. The complete details of these Experiments 2-4 are reported in the Supplementary Information Text (SI Text) 5.

Effort level was manipulated block-wise (Fig. 1, Experiment 1, 4-minute blocks). In Experiment 1 the cognitive effort variant required performing trials of the Multi-Source Interference Task (MSIT, Fig. 2, Bush & Shin, 2006). The high effort condition required completing interference trials (demanding more cognitive effort), and the low effort condition required completing congruent trials (demanding relatively less cognitive effort). The physical variant of the task required participants to rapidly press a key to reach a new patch (Fig. 2 right panel, based on previous research demonstrating that rapid key-pressing is physically effortful and costly, (Treadway et al., 2009)). The high effort condition required participants to press the key the maximum number of times they could in the time allotted (individually determined in a preceding calibration phase), and the low effort condition required half that number of key-presses. Travel time (i.e., time to complete the MSIT or key-pressing tasks) was fixed and the same across both variants of the task and the high and low effort conditions of each. We predicted effort-avoiding participants would have a lower patch leaving threshold (in units of apples) in the high effort conditions compared to the low effort conditions, since travel (effort) costs were greater in the former. The measure of effort cost for an individual was the differential travel cost of the more effortful condition (incongruent MSIT, or Larger Number of Presses) compared to the less effortful condition (congruent MSIT, or Smaller Number of Presses).

**Figure 1:**
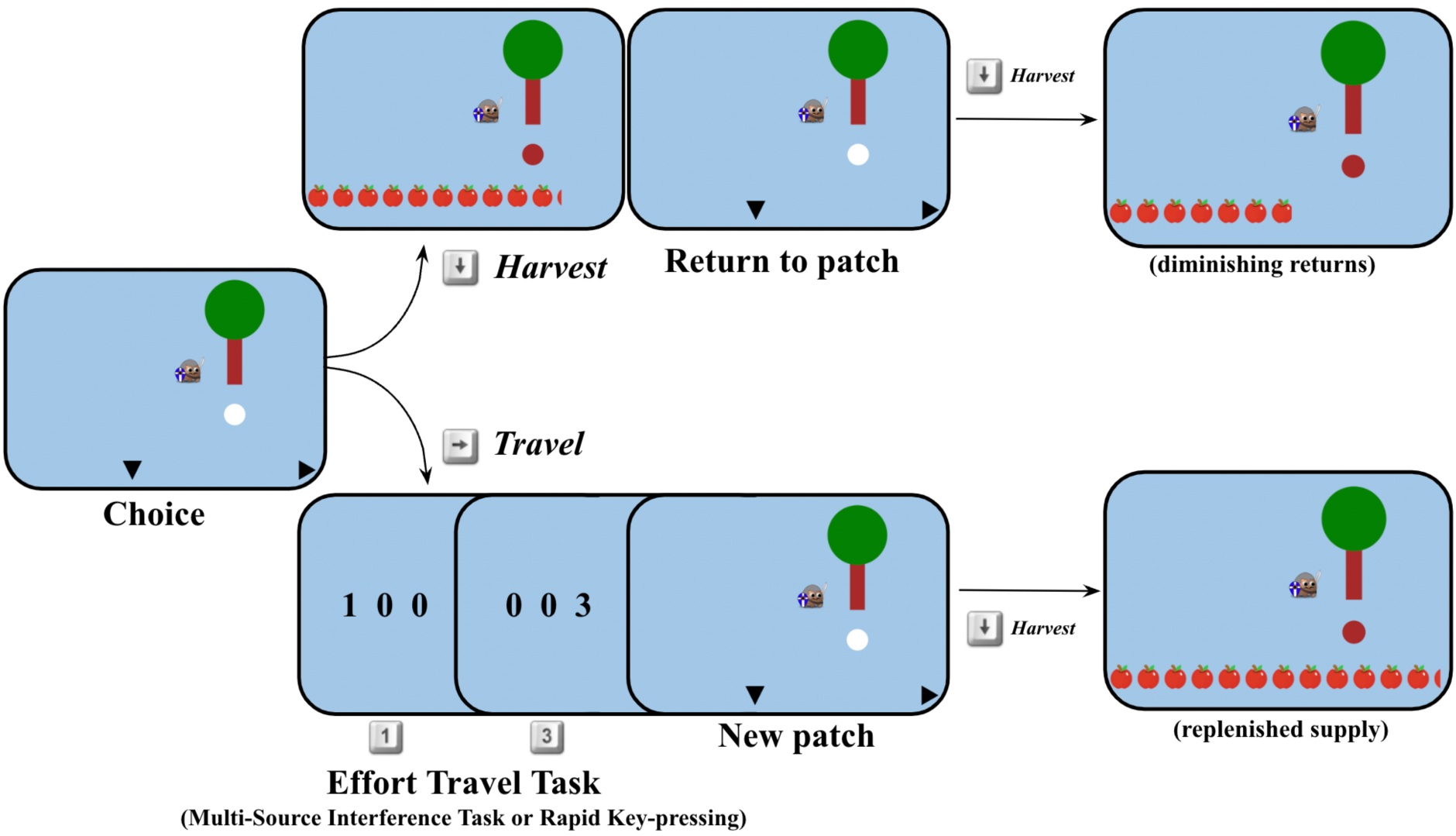
Foraging trial diagram. On each trial participants chose to harvest the tree they were at (down arrow key) or travel to a new tree (right arrow key), during the travel they completed an effortful task, after which they arrived at a new patch with a replenished supply of apples.

**Figure 2:**
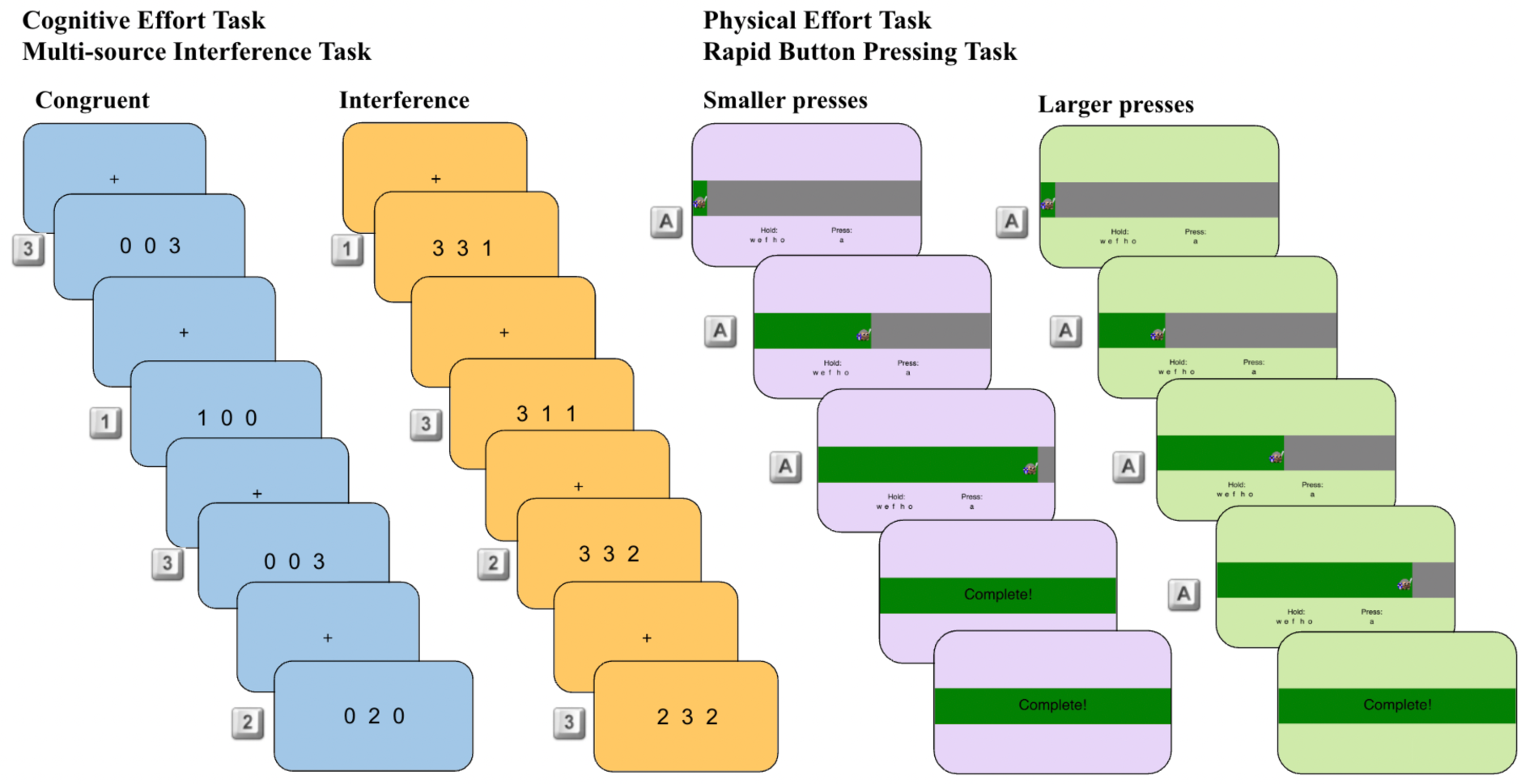
Effort travel tasks Experiment 1. Left panel: Cognitive Effort, Multi-source Interference Task. Participants identified which number was the oddball in a list of three numbers. The background color differed for the high effort (interference trials, orange) and low effort (congruent trials, blue) conditions. They responded with the ‘1’, ‘2’, ‘3’ keys. Interference trials have a competing distractor response for which the oddball target is flanked by the distractor and in the spatial position of the distractor. The correct response for each example screen is displayed on the left of that example screen. Right panel: Physical Effort, Rapid Key-pressing Task. Participants rapidly pressed the ‘a’ key while holding down the ‘w’, ‘e’, ‘f’, ‘h’ and ‘o’ keys. Pressing the ‘a’ key moved the avatar rightwards and filled up the grey horizontal bar with green. When participants reached the goal number of presses ‘Complete!’ appeared in the horizontal bar and participants waited for the remainder of the travel time. The background color differed for the high effort (smaller presses, purple) and low effort (larger presses, green) conditions.

#### Advantages compared to previous tasks

Our motivation to develop the Effort Foraging Task was to provide a novel approach to effort cost estimation that may help improve predictive validity. In particular, relating the behavior observed in effort-based decision tasks to external measures such as self-reports has been a challenge (Juvina et al., 2018; Eisenberg et al., 2019; Strobel et al., 2020).

One potential problem is that existing tasks have involved hypothetical choices, or choices separated in time from a later realization of cognitive effort (Westbrook et al., 2013; Lopez-Gamundi & Wardle, 2018). In the Effort Foraging Task participants experience the effort directly and immediately, after each choice to travel. Furthermore, any previously reported tasks have involved explicit effort-based decision making; for example, participants are asked directly if they would rather complete a high or low effort task (Kool et al., 2010; Westbrook et al., 2013; Lopez-Gamundi & Wardle, 2018). Explicit decision making may be subject to secondary demand characteristics, in which participants behave according to what they think the experimenter wants or with their self-image. Decisions that explicitly trade off numeric quantities are also susceptible to idiosyncratic arithmetic heuristics (Marzilli Ericson et al., 2015). In addition, direct tasks, such as the Cognitive Effort Discounting Paradigm (Westbrook et al., 2013) or the Effort Expenditure for Rewards Task (Treadway et al., 2009) may engage real-world economic considerations (e.g., that one should be paid more to work more), which may obscure or interfere with effort seeking behaviors that may otherwise occur in naturalistic settings. This is consistent with the observation that effort seeking is not reported in direct tasks (i.e., reverse effort discounting, preferring to do a more demanding task for less money over a less demanding task for more money Treadway et al., 2009; Westbrook et al., 2013). In contrast, the influence of effort in the Effort Foraging Task is measured indirectly; participants learn about the environment reward rate and effort costs through experience, and their choices reflect their ongoing evaluation of these quantities. Interestingly, we consistently found a subset of participants who appeared to be effort-*seeking* in this task, in line with the idea that secondary demand characteristics obscure or interfere with effort seeking behavior in explicit tasks.

Another concern common to previous studies is the presentation of two options simultaneously, as this may distort choices or complicate their interpretation (Kool et al., 2010; Westbrook et al., 2013; Lopez-Gamundi & Wardle, 2018). Research in intertemporal choice has shown that rodents and primates (including humans) are less impulsive decision makers when they are making serial rather than simultaneous choices. It has been hypothesized that this is because serial decision making is more ecologically valid (evaluating each single option in isolation and choosing to accept or reject it and search for another, Blanchard & Hayden, 2015; Carter et al., 2015; Carter & Redish, 2016). The Effort Foraging Task is serial, as participants decide to either accept the current harvest value or reject it and travel to a new patch. Formal analysis of these foraging decisions using the Marginal Value Theorem provides a theoretically motivated and quantitatively rigorous approach to measuring effort costs.

## Results

The primary dependent variable in our analyses were exit thresholds (the expected reward for harvesting when participants’ chose to travel), which reflect the point when the cost of leaving just offsets the benefits of reaching a replenished patch (which is increasing as the current patch yields progressively less). In line with the Marginal Value Theorem (1) we assumed that participants set their exit thresholds to maximize the rate of rewards minus costs per time step. As all of the reward rate variables are observed, this allows us to solve for the subjective cost *c* that best rationalizes the observed exit behavior. We implemented a computational model based on the Marginal Value Theorem and fit it to participants’ exit thresholds to quantify the relative increase in travel cost between the high and low effort conditions for each effort type (Analysis methods, ‘Hierarchical Bayesian Marginal Value Theorem Model’).

### Summary of results

We found that differences in foraging decisions (viz., exit thresholds) are a useful indirect measure of motivation to exert both cognitive and physical effort. Consistent with our prediction, average exit threshold was lower in the higher travel effort than the lower travel effort conditions (Fig. 3). Participants (Experiment 1, N=537) opted to stay longer in a patch, accepting diminishing rewards, in the high travel effort conditions to avoid the increased cost of travel. Results from Experiments 2-4 confirmed that participants’ behavior remains similar across manipulations of cognitive effort type (Experiment 2 (N-Back)), environment richness (Experiment 3 (Richness)), and with explicit travel time instructions (Experiment 4). Fits of the Marginal Value Theorem model to trial-by-trial behavior (Experiment 1 (MSIT)) further confirmed the presence of high effort costs and revealed an interesting mixture of effort-avoiding and effort-seeking participants. We found that cognitive and physical effort costs were moderately positively correlated, and that both effort costs had patterns of correlation to self-report measures related to motivation and affect.

**Figure 3:**
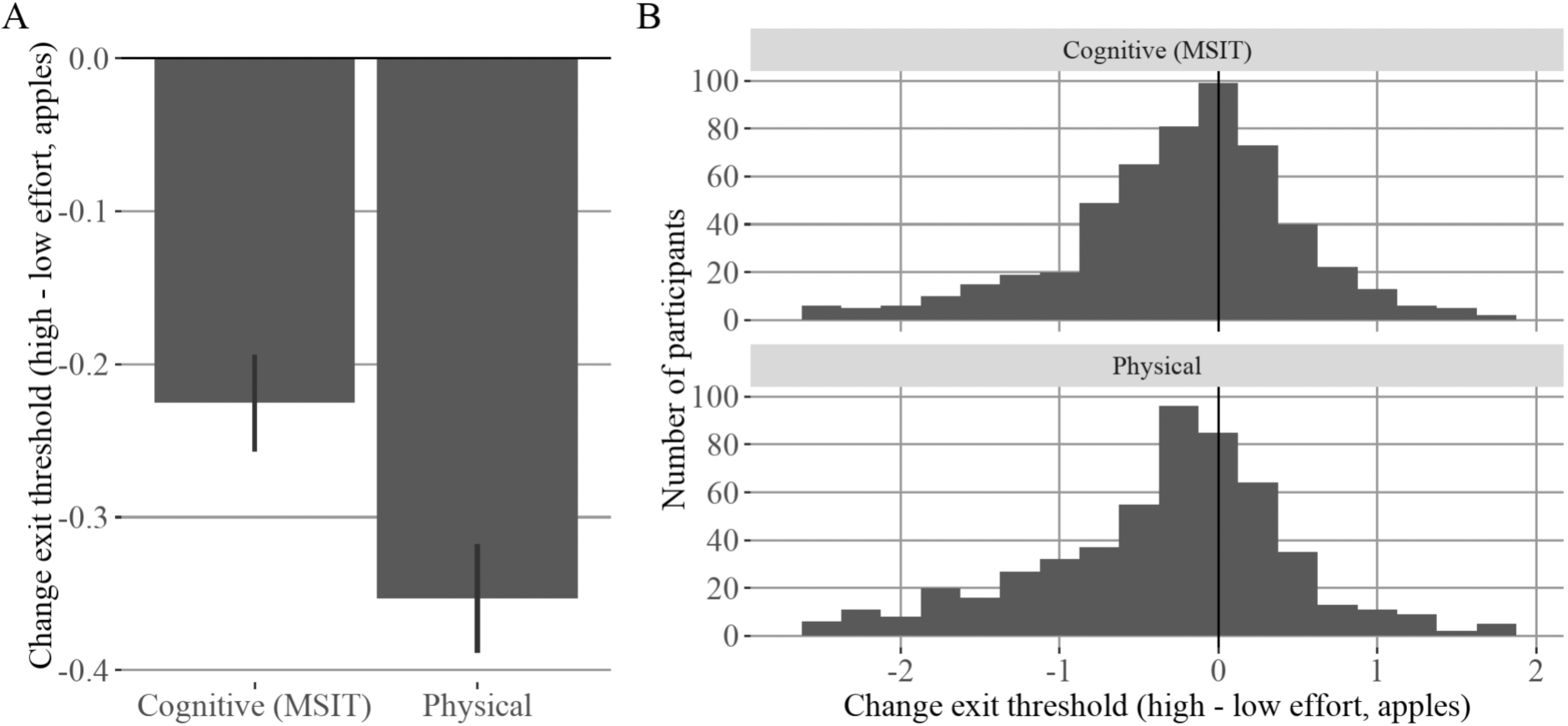
Change in exit thresholds by effort condition. Experiment 1 (MSIT). A: y-axis: Group-level mean change in exit threshold for cognitive and physical effort. x-axis: effort type. As predicted, on average participants exhibited lower exit thresholds in the high relative to low effort conditions. Error bars indicate standard error of the mean. B: Individual variation in change in exit threshold. Top row: Histogram of participants mean change in exit threshold for cognitive high effort relative to cognitive low effort. Bottom row: mean change in exit threshold for physical high effort relative to physical low effort. Most participants were effort-avoiding (negative change in threshold), whereas some participants showed indifference to effort condition (near zero) or were effort-seeking (positive change in threshold).

### Change in exit threshold by effort condition

As a model-agnostic metric of high effort cost, we used the change in exit threshold from low to high effort conditions. For each participant we computed the average exit threshold per condition (see overall threshold results in Fig. S1 and Fig. S2), and the difference between them (high effort - low effort mean threshold). We expected this value to be negative, reflecting effort avoidance. If threshold increased for a participant, this suggested effort seeking. Across participants, we found a mix of effort avoidance, effort seeking, and indifference to effort (values close to zero) (Fig. 3 right panel). We computed the group average change in threshold (Fig. 3) and used linear mixed-effects regression to test whether change in exit thresholds significantly differed from zero. As predicted, on average, participants exited trees later in the high relative to low effort conditions (mixed-effects regression: interference - congruent MSIT, *β_cognitive_* = −0.236 apples, df=460.071, F=50.062, p<0.001, Larger - Smaller Number of Presses *β_physical_* = −0.379 apples, df=474.041, F=87.326, p<0.001). We estimated the split-half reliability of the change in exit threshold measure, which was r=0.85 for cognitive effort, and 0.82 for physical effort (details in SI Text 2). As a model-agnostic estimate of the relationship between cognitive and physical effort cost we measured the correlation of the change in exit threshold across cognitive vs. physical effort conditions and found a significant positive correlation (r=0.203, t=4.78, df=535, p<0.001). Next, we used the foraging behavior to formally quantify the additional cost of the high effort tasks using a model based on the Marginal Value Theorem.

### Hierarchical Bayesian Marginal Value Theorem model to estimate effort costs for an individual

We fit a hierarchical Bayesian logistic model based on the Marginal Value Theorem (Charnov, 1976), which predicted harvest versus exit decisions by comparing expected reward on the next harvest against the average reward rate (see Analysis methods, ‘Hierarchical Bayesian Marginal Value Theorem Model’). We defined the reward rate in terms of known reward rate values of the foraging environment per effort condition per participant (apples earned, time cost incurred, number of patches visited), as well as the unknown reward rate value (the cost of travel). The cost of travel in high effort blocks was expressed as the marginal increase in cost of travel from low to high effort. Defining this cost as a difference measure controls for any additional biases individual participants may have (such as differences in the subjective value of the reward) which are common to both conditions. The dependent individual differences measures in this task were the inferred cognitive and physical effort cost parameters. The other model parameters were the travel costs in the cognitive and physical low effort conditions, and the inverse temperature applied to the softmax function (higher values indicate less noisy effects of rewards and thresholds on choices).

Consistent with the model-agnostic change in threshold metric, the group-level posterior parameter fit indicated that both high-effort tasks were costly on average (Tab. S1). There was a range of individual differences (Fig. 4), cost was positive for most participants, some participants were indifferent to the effort manipulation (cost near zero), and some participants had a negative cost (cognitive, N=78, 14.5% of sample, physical, N=67, 12.5% of sample), suggesting that effort was valued.

**Figure 4:**
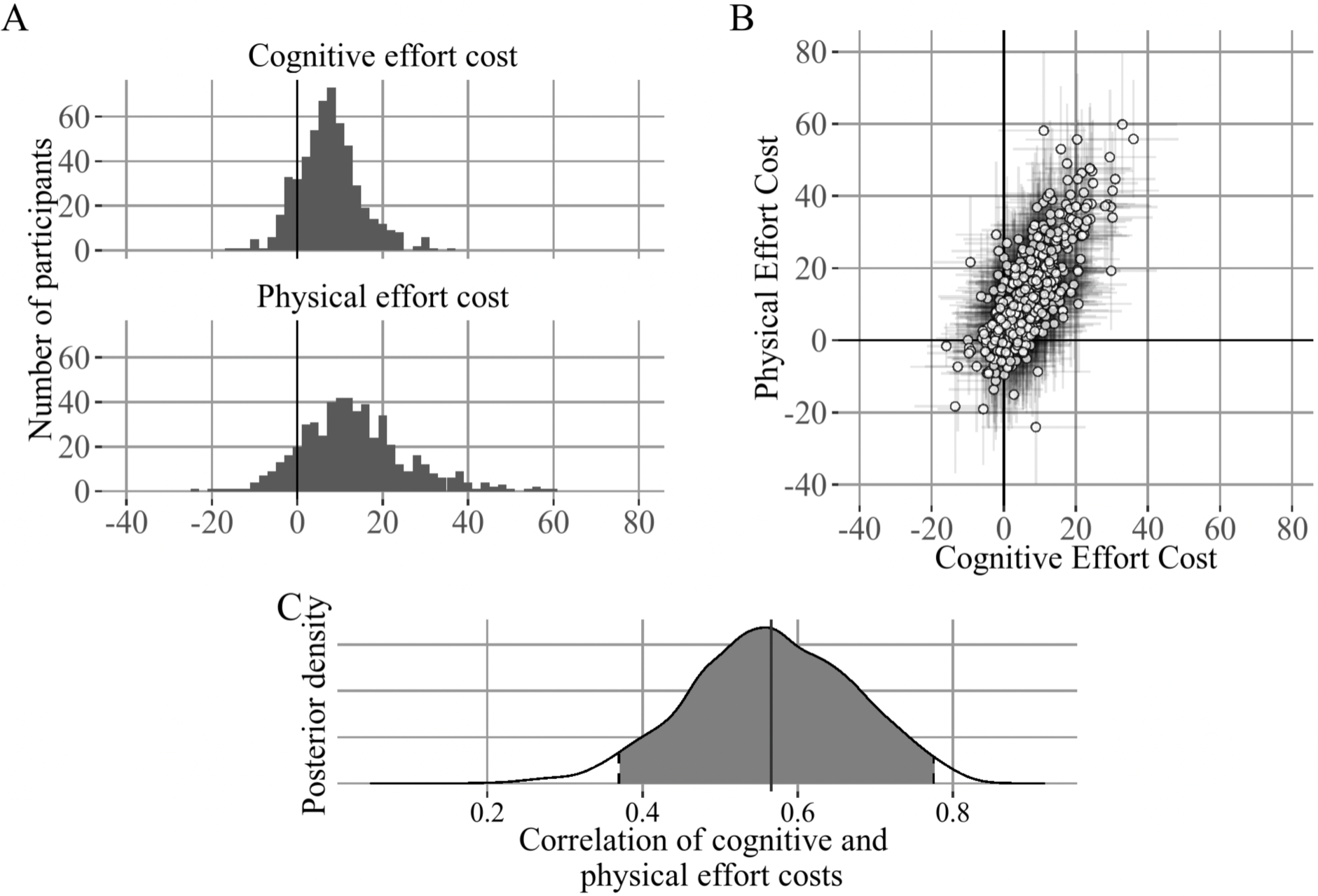
Correlation between individual differences in cognitive and physical effort costs. Experiment 1 (MSIT), A: Individual differences in the high effort travel costs (expressed as the additional cost of the high relative to the low effort condition). Paralleling the pattern of exit thresholds most participants experienced the high effort conditions as effortful (positive cost), whereas some participants were insensitive to the effort manipulation (cost near zero) and others were effort-seeking (negative cost). B: x-axis: Individual differences in cognitive effort costs, y-axis: Individual differences in physical effort costs. Error bars indicate 80% HDI. C: Posterior distribution of correlation between high effort cost for cognitive and physical effort. Cognitive and physical effort costs are positively correlated (correlation=0.566, 95% HDI=0.355 - 0.766).

### Relationship between cognitive and physical effort costs

We estimated the correlation across participants between the cognitive and physical high effort costs (again, each estimated as reflecting the additional cost of high effort relative to the low baseline) using the participant-level covariance matrix when fitting the MVT model. We found a moderate positive relationship (mean correlation=0.566, 95% HDI=0.355 - 0.766, Fig. 4). This suggests a potential common representation for costs of different types used in effort-based decision making. We confirmed this correlation was not driven by the subset of participants who showed negative effort costs by re-fitting the model omitting participants with negative cognitive or physical effort cost (Fig. S3).

### Travel task performance relationship to effort costs

For cognitive effort, we tested whether cognitive task performance contributed to the cognitive effort cost measured by foraging choices (see individual differences in MSIT performance in Fig. S4 and relationship between cognitive and physical effort costs and travel task performance in Fig. S5). We regressed the difference in (log transformed) MSIT error rate and (log transformed) reaction time onto cognitive effort costs. We found that the difference in error rate significantly predicted cognitive effort cost (estimate=15.313, SE=3.684, t=4.156, p<0.001), indicating that participants with higher costs performed worse on the MSIT. However, the reaction time interference effect did not predict cognitive effort cost (estimate=4.861, SE=3.802, t=1.278, p=0.202). In this regression the intercept was not significantly different from zero, suggesting that the effort costs measured are performance-related (estimate=4.861, SE=3.802, t=1.278, p=0.202, compared to an intercept-only model estimate=7.548, SE=0.323, t=23.38, p<0.001). We see the same qualitative result using robust regression. This finding suggests participants may adaptively calibrate their effort costs according to their error rates, and/or that effort costs and error rates are both affected by more general task engagement or motivation (see Discussion).

Next, we investigated the analogous relationships for physical effort. We regressed two measures of performance, the percentage of uncompleted presses in the smaller and larger presses condition, and the required number of presses determined in the calibration phase, onto physical effort costs. We found no relationship with physical effort costs and the percentage of uncompleted presses in the smaller (estimate=-0.387, SE=0.234, t=-1.656, p=0.098) nor the larger (estimate=0.112, SE=0.140, t=0.802, p=0.423) presses condition, nor to the required presses (estimate=0.018, SE=0.088, t=0.198, p=0.843). The physical effort cost effect remained controlling for all these variables (intercept estimate=13.20, SE=2.08, t=6.33, p<0.001). One potentially important difference is that the physical effort requirement was individually calibrated such that completion rates were near ceiling (Fig. S4). Lastly, we correlated the error rates across the high-effort cognitive and physical effort tasks and found a weak positive correlation (r=0.15, t=3.52, df=535, p<0.001).

### Relationship to self-reported motivation and affect

We conducted an exploratory canonical correlation analysis (CCA), to examine the relationships between our task measures and self-report surveys. The high dimensionality of both the survey and task measures poses a multiple comparisons issue for correlation or regression analyses. In previous work, we and others have used dimensionality reduction techniques such as factor analysis to summarize key dimensions of survey data prior to regressing them on individual task measures (Gillan et al., 2016). Here, we build on that approach by using CCA, a dimensionality reduction technique that simultaneously performs dimension reduction on two domains of data, to identify summaries of each domain that maximally relate to one another (here, the relationship between surveys and task behavior measures, Wang et al., 2020). This allowed us to both examine the external validity of our measures and to investigate the broader context associated with effort sensitivity (with data from N=430 Experiment 1 participants who completed the survey and passed attention checks, see Fig. S6, details in Analysis methods, ‘Canonical correlation analysis.’). The dependent variables (N=11) were all the self-report summary scores, (see Tab. S2). The predictor variables (N=5) were all the task behavior measures of interest: (log transformed) error rate on congruent and incongruent trials, cognitive and physical effort costs, and overall threshold. Including multiple task behavior measures allowed us to explore the structure of multivariate relationships between (potentially partially correlated) task measures such as effort costs and task performance, and on the other hand, several proxies of real-world behavior. This approach has the benefit of increasing sensitivity (by making use of all the measures simultaneously), while reducing the risks of multiple comparisons (by treating all of these factors in a single omnibus analysis).

CCA revealed five dimensions, of which one was significant (summarized in Tab. 1, and full result shown in Fig. S7, Wilks’ Lamda (Wilks, 1935), using F-approximation, dimension 1: correlation=0.283,stat=0.819, F-approx=1.538, df1=55, df2=1919.899, p<0.0072, dimension 2: correlation=0.257, stat=0.891, F-approx=1.222, df1=40, df2=1575.487, p<0.162, dimension 3: 0.161, dimension 4: 0.130, dimension 5: 0.063, p>0.83 for dimensions 3 to 5). To interpret which task behavior and/or self-report measures contributed most strongly to the significant dimension, we highlighted those that had a coefficient greater than 0.5 along each dimension (Tab. 1). On the task measure side, the first dimension was most closely associated with increased MSIT Error Rates and decreased Cognitive Effort Cost. On the survey side, this dimension was most closely associated with decreased (self-reported) Cognitive Function, increased Behavioral Activation, decreased Anxiety, and decreased Self-efficacy. Overall, Dimension 1 may capture a pattern of individual differences ranging from cautious/error-averse attentiveness to error-prone inattentiveness to the tasks (similar to Moutoussis et al., 2021). This is reflected (on the self-report side) in a spectrum from high anxiety to high behavioral activation, which is related on the task side with increasing error rates. Intriguingly, this dimension is also associated with decreased cognitive effort costs (but, conversely, increased physical effort costs, Fig. S7), a dissociation that speaks against the idea that this type of task engagement drives the positive univariate correlation between cognitive and physical effort costs.

**Table 1:** Canonical Correlation Analysis Result Summary Experiment 1 (MSIT). Column 1, Canonical Dimension, correlation, and p-values. For each dimension coefficients larger than 0.5 are displayed in column 2 (for task behavior variables) and in column 3 (for self-reports). Arrows indicate positive or negative coefficients. Dimension 1 was significant whereas Dimension 2 was not. Dimension 1 reflected increased MSIT Error Rates and decreased Cognitive Effort Cost, whereas Dimension 2 reflected decreased Physical Effort Cost, suggesting each of these factors have a distinct relationship with respect to external correlates. The associated self-report coefficients in Dimension 1 were decreased Cognitive Function, increased Behavioral Activation, decreased Anxiety, and decreased Self-efficacy.

| Canonical Dimension (correlation, *p-value*) | Task behavior (coefficient) | Self-reports (coefficient) |
| --- | --- | --- |
| **Dimension 1** (0.283, 0.007) | ↑Congruent Error Rate (0.66) | ↓Cognitive Function (-0.75) |
|  | ↑Interference Error Rate (0.58) | ↑Behavioral Activation (0.74) |
|  | ↓Cognitive Effort Cost (-0.52) | ↓Anxiety (-0.62) |
|  |  | ↓Self-efficacy (-0.51) |
| **Dimension 2** (0.257, 0.162) | ↓Physical Effort Cost (-0.92) | ↑Anxiety (0.73) |
|  |  | ↓Need for Cognition (-0.71) |
|  |  | ↓Fatigue (-0.63) |

Dimension 2 was not significant, though the numerical correlation was comparable (canonical correlation=0.257, p>0.162) to Dimension 1. We inspected it to visualize what types of structure (albeit non-significant) remain residually in the data after the removal of Dimension 1. Dimension 2 was more closely related, among task measures, to decreased Physical Effort Cost, and was associated, among self-reports, with increased Anxiety, decreased Need for Cognition, and decreased physical Fatigue. Thus, although not significant, Dimension 2 may reflect a plausible relationship, which could be sought in future, higher-powered studies, between physical fatigue and physical effort cost, as distinct from cognitive effort cost.

### Validation experiments

In addition to the main experiment (Experiment 1 (MSIT)), we tested three other versions (Experiments 2, 3, and 4) to validate key findings (see SI Text 5).

#### Generalizability of cognitive effort manipulation

In Experiment 2 (N-Back), we tested a different form of effort as the travel manipulation (N=81 included, see exclusions, quantitative cutoffs, and number of outlier participants in Tab. S3). Specifically, we compared foraging behavior when the travel task was the 3-Back versus 1-Back level of the N-Back task (Nystrom et al., 2000). As predicted, on average across participants exited trees later in the high (3-Back) relative to low (1-Back) cognitive effort conditions; linear mixed-effects regression estimate for N-Back (3-Back - 1-Back): = −0.504 apples, df = 75.981, F = 30.339, p < 0.001, physical (smaller-larger) = −0.448, df = 75.170, F = 27.151, p < 0.001. The MVT model fit also indicated a positive high effort cost (Tab. S4). Although much smaller in size, Experiment 2 did not replicate the correlation observed in Experiment 1 between cognitive and physical effort cost as estimated by the MVT model (see Fig. S8). The highest density interval (HDI) was very wide (−0.38 to 0.45) suggesting Experiment 2 (N=81) may have been underpowered to detect a correlation similar in size to that seen in Experiment 1. Another possibility is that the size of any correlation may have been reduced by a selection bias in Experiment 2 (i.e., admission to university, Herńan & Monge, 2023, SI Text 5.A.4.).

#### Collateral predictions of the Marginal Value Theorem

In Experiment 3 (Richness), we tested adherence to classic predictions of the Marginal Value Theorem, not tested in Experiment 1, by manipulating patch richness. Leaner environments (yielding lower overall mean reward rate) should be associated with lower exit thresholds relative to richer environments. To test for this effect, we compared two levels of reward richness, by adjusting the mean of a normal distribution used to draw the initial yield of a patch. As predicted, we found that participants lowered their exit thresholds in the lean compared to rich conditions (Richness contrast; sum sq. 0.788, mean sq. 0.788, DenDF=27.95, F=10.49, p<0.0031).

#### Explicit instruction travel time is fixed

Research shows that time perception is subjective, and that a more demanding task can make subjective time estimates more variable and less accurate (R. A. Block et al., 2010). If subjective travel time estimates were generally longer in the high effort condition, that could contribute to effort avoidance (though higher cognitive load can also shorten subjective time estimates, Khan et al., 2006; F. Block & Gellersen, 2010). In Experiments 1, 2, and 3, we did not instruct participants that the travel time was fixed regardless of the effort condition, leaving open the possibility that participants’ subjective time estimates differed between travel task conditions, and contributed to the observed effect. In Experiment 4 we evaluated this possibility by including explicit instructions that the travel time was fixed across all conditions. The results replicated the findings of effort avoidance in Experiment 1. As predicted, on average participants still exited trees later in the high relative to low effort conditions for both cognitive and physical effort (linear mixed-effects regression estimate for MSIT (Interference-Congruent):= −0.318 apples, df=49.50, F=12.66, p<0.001, physical (smaller-larger) = −0.391, df=47.72, F=5.66, p<0.021, see Fig. S9).

## Discussion

We developed the Effort Foraging Task to quantify the costs of cognitive and physical effort at the level of the individual. Participants played a computer game in which they could forage for virtual apples in a patch with diminishing returns or abandon that patch for a new (initially) richer patch at the expense of time and effort. Participants completed blocks of the task in which the travel cost was either cognitive or physical effort, each at one of two difficulty levels (high and low effort). We measured their ‘exit threshold’ as the number of apples the participant could have expected to get on their next harvest on trials in which they decided to travel instead. We found that on average participants lowered their exit threshold (staying longer, accepting diminishing returns) in the high relative to low effort conditions, consistent with the high effort task having a monetary cost.

Further analyses in Experiment 1 demonstrated that these cognitive effort costs are correlated with differences in error rates between the low and high cognitive effort tasks, suggesting that the costs may at least partially reflect error avoidance, and/or that effort costs and error rates are both affected by more general task engagement. Expected Value of Control model simulations (Musslick et al., 2018) demonstrated the problem of identifiability of effort costs versus other factors that contribute to cognitive effort allocation: skill and reward sensitivity. That is, if someone avoids effort (i.e., restricts allocation of cognitive effort to a demanding task) this could reflect a higher cost of effort, but it could also reflect poorer ability and/or weaker incentives. These individual differences would impact choices in the foraging task, and impact or otherwise index performance on the cognitive effort travel task. To address these potential confounds, we employed a multidimensional, canonical correlation analysis, offer a richer view on the interrelationship between multiple variables.

### Relationship to self-reported motivation and affect

CCA revealed inter-relationships between Effort Foraging Task variables and self-report proxies of real-world motivation and psychiatric symptoms such as cognitive function, anxiety, behavioral activation, and self-efficacy. One significant dimension was identified that loaded on multiple cognitive task variables and self-report measures (Tab. 1). This dimension plausibly captures something like cautious or compliant attentiveness to the tasks: specifically, on the task side, it reflects increased error rates, which, on the self-report side, are correlated with movement along a spectrum from anxiety to behavioral activation. However, this dimension also dissociates cognitive effort sensitivity (which is decreasing in this dimension) from both error rates and physical effort sensitivity (which are increasing). These dissociations also argue against the possibility that variation in this type of task engagement (Moutoussis et al., 2021) drives the univariate correlation between cognitive and physical effort costs. Altogether, Dimension 1 suggests a more complex profile than simply nonspecific attentional engagement. These rather nuanced results may be enabled by our relatively careful exclusion of grossly inattentive participants (both through attention check items in the survey and task behavior-based exclusions), who can otherwise drive uninformative correlations and obscure more informative relationships (Zorowitz et al., 2021).

Sensibly, we found that individuals who reported better cognitive function in the past week (and/or reported fewer cognitive difficulties on the survey), and higher self-efficacy, exhibited better performance in the cognitive effort travel tasks in Dimension 1. However, the positive association between such cognitive function, self-efficacy, and cognitive effort cost in Dimension 1 is contrary to our predictions. It may be that the subjective evaluation of cognitive function and self-efficacy are more related to error proneness than to cognitive effort cost. Alternatively, it may be that participants who are experiencing worse cognitive function and lower self-efficacy are less sensitive to the cognitive effort task demand differences. Lastly, while we did not have any *a priori* hypotheses about anxiety’s relationship to effort costs, it figures strongly in predicting both effort costs and error rates in Dimension 1, meriting further investigation into how effort costs might relate to symptoms of anxiety. The CCA is an exploratory analysis and factor structure identified should be replicated in a confirmatory sample in future work. Informed by these results, future work could go beyond self-report, and use ecological momentary assessment measures (Strobel et al., 2020; Krönke et al., 2021; Crawford et al., 2023).

### Cognitive and physical effort relationship

Our design measures cognitive and physical effort costs on a common scale, revealing a significant and substantial correlation between them. This suggests that a common mechanism may compute costs across multiple domains, consistent with research showing that; i) overlapping in brain areas are involved in cognitive and physical effort decisions using human neuroimaging (Schmidt et al., 2012; Chong et al., 2017), ii) intermixing choices about cognitive and physical effort affects choices for both effort types (Toro-Serey et al., 2021), iii) cognitive fatigue impacts physical effort exertion and vice versa (Marcora et al., 2009; Giboin & Wolff, 2019).

Indeed, one set of mechanisms that may be shared between the domains in our study, and contribute to their relationship, is the cognitive and neural mechanisms for foraging decisions themselves. Spatial foraging is hypothesized to be an evolutionary antecedent of abstract cognitive search processes such as memory search (Hills et al., 2008). As such, shared processes for foraging (both external and mental) have been proposed to be a core component of cognition (Pirolli & Card, 1999; Wilke et al., 2009; Hills et al., 2012; Todd et al., 2012; Wolfe, 2013; Hills et al., 2015). Hills, Todd, and Goldstone (2008) demonstrated, for instance, a causal connection between spatial and mental foraging; they manipulated the distribution of resources in a spatial foraging task and observed that this affected behavior on a subsequent mental foraging task (involving a word search puzzle). This interplay between the cognitive and physical domains offers an intriguing explanation for the observed relationship between individual differences in cognitive and physical effort costs, and an interesting set of hypotheses about interaction across the domains to pursue in future work with the Effort Foraging Task.

Foraging provides an ideal framework to understand interactions between action costs (e.g., cognitive and physical effort costs, time costs, and factors such as risk and uncertainty, Kilpatrick et al., 2021) as they are all at play in naturalistic foraging behaviors. For example, exerting greater physical effort (e.g., vigor, Niv et al., 2007) can reduce time costs, and exerting greater cognitive effort (i.e., planning an efficient action) can reduce both physical effort and time costs. The present study moves in this direction by adding a cognitive or physical effort task requirement to travel in a virtual patch foraging environment with human participants. To directly test one aspect of interplay between cognitive and physical effort decisions, the task could be adapted to intermix cognitive and physical blocks. To ascertain the effect exposure to one domain on decision making has on the other, a study could compare effort type order effects (cognitive before physical versus physical before cognitive), as dissociated from the effect of overall experiment time and fatigue (double length cognitive, and double length physical).

### Generalizability

The Effort Foraging Task, as well as existing alternatives, are constrained to a limited set of pragmatic laboratory tasks which may not be representative of the putative broader domains of cognitive and physical effort. Experiment 2 offers some evidence for the generality of our approach across distinct operationalizations of cognitive effort, since it produced broadly similar results (e.g., predominantly effort avoidance, some effort-seeking) using a different putatively cognitively effortful task (MSIT vs N-Back) than Experiment 1. However, the extent to which this holds across the much wider range of tasks that can be considered “physical” or “cognitive” (Neisser, 1967), and whether such different tasks track each other in terms such as individual differences or external correlations in effort avoidance, remains an important question for future research. Within-participant comparisons using a broad range of cognitive and physical travel tasks could speak to the domain generality of cognitive versus physical effort costs. Intriguingly, research using the Cognitive Effort Discounting Paradigm found correlations between effort avoidance for two distinct cognitive control domains (working memory and speech comprehension during background noise) suggesting a task general component of cognitive effort costs (Crawford et al., 2022). Longitudinal applications of this task could be used to characterize trait-like versus state-dependent contributions to behavior. State variables such as time of day, arousal level, hunger, affect, the presence of psychiatric symptoms, and others may all contribute to effort-based decision making. Understanding state-dependent influences could prove valuable in identifying tractable tools to promote effort exertion in daily life. Relatedly, test-retest reliability should be measured in a replication study; here, we found the split-half reliability to be above 0.8 for both effort costs.

### Effort Seeking

In all the experiments reported here, we consistently observed a subset of participants who exhibited *negative* effort costs (i.e., a preference for the high effort option over the low effort option) for both the cognitive and physical effort conditions, suggestive of effort seeking. Both cognitive and physical effort seeking occur frequently in real world behaviors and have been linked in the psychological literature to “need for cognition” and “learned industriousness” among other constructs. Evidence shows instances in which effort requirements *add* value to a reward (as opposed to discounting rewards, reviewed in Inzlicht et al., 2018). Although this comprised a minority of participants in our experiments (14.5% cognitive, 12.5% physical in Experiment 1), it nevertheless suggests the need for extensions to existing utility models (e.g., the Expected Value of Control framework). Cognitive effort seeking, *prima facie*, indicates positive value assigned to exerting cognitive effort, which may reflect — directly or indirectly — longer term value attached to information-seeking and learning that yield better future performance (Geana et al., 2016; Agrawal et al., 2019). In the same vein, physical effort seeking may reflect value attached to future performance improvements, for example via physical learning (e.g., skill acquisition, strength building), and information seeking in physical space. In both cases effort seeking may also have to do with boredom, which may hold a disutility that encourages application of effort (e.g., Agrawal et al., 2022). This can be adaptive because effort expenditure can help to obtain rewards, in which case doing nothing carries an opportunity cost. Individual differences in the value assigned to novel information/learning (e.g., information bonus, Wilson et al., 2014) may also have contributed to the behavior of participants with negative effort costs, for whom the information/learning value would be higher in the high effort conditions, leading to a preference for the high effort conditions. Each of these factors likely comprise their own dimensions of individual variation (e.g., boredom aversion, information bonuses) that were not measured in our experiments. An additional variable which might contribute to effort avoidance versus effort seeking is distortions in subjective time perception (Khan et al., 2006; F. Block & Gellersen, 2010; R. A. Block et al., 2010). Specifically, if subjective times in the low effort conditions were exaggerated for some participants (e.g., if for them, greater engagement leads to a perception that less time has passed), this could lead them to perceive a net increase in the travel cost relative to the high effort conditions, promoting effort seeking. However, in Experiment 4 we confirmed that the main effect of effort avoidance was conserved even when participants were explicitly instructed that the travel time was fixed. Future research is needed to further investigate the factors that drive effort seeking in this task and others.

### Testing effort cost theories

Further work using the Effort Foraging Task may be useful in testing alternative accounts of the basis of effort costs (i.e., opportunity cost, processing, and metabolic accounts Baumeister & Heatherton, 1996; Kurzban et al., 2013; Musslick & Cohen, 2021). For example, to test opportunity cost accounts, an experiment could manipulate whether a low effort alternative task is available (e.g., browsing social media instead of completing the Effort Foraging Task for money). Opportunity cost accounts would predict that the cognitive effort cost measured by foraging behavior would be higher during periods in which an alternative was on offer (Kurzban et al., 2013). To test cost of processing accounts, the travel task could involve multi-tasking. By these accounts, participants should treat multi-task sets that recruit more shared representations as more costly than sets that recruit more separated representations (Musslick & Cohen, 2021).

Effort-based decision making has considerable importance in daily life. Critical questions remain about how to disentangle aspects of motivation for effort, how these aspects are represented in the brain, and the role they play in real-world behaviors. The cognitive computational study of motivation has the potential to help people reach their goals by identifying the mechanisms of motivation and ways to enhance motivation towards what matters most to an individual.

## 2 Methods

The Effort Foraging Task adapts a version of the patch foraging paradigm by embedding cognitive and physical effort costs in between patches (here, a simulated orchard with apple trees). The logic of the task is described in the Introduction section ‘The Effort Foraging Task’ and complete details on the task and analysis are described below. On each trial an image of a tree appeared on the screen, representing an immediately available source of reward. Participants could choose to harvest that patch (tree) or travel to a new, replenished patch (Fig. 1). When a tree was harvested it ‘shook’ and apples were displayed under it (apples were displayed in a single, left justified, row). Reward depleted within a patch such that the more times a tree was harvested the fewer apples it produced. When participants choose to exit the patch, they had to “travel” which consisted of completing a cognitively or physically effortful task. Participants had a fixed amount of time to collect apples (money). Therefore, they must balance the diminishing returns associated with staying at a patch with the travel costs required to reach a new, replenished patch.

To indicate the start of a trial a circle below the tree turned white and participants were able to make their decision. The circle below the tree was brown when participants could not enter a decision (apples being displayed and waiting through harvest delay). If participants took too long to decide (1 second deadline) a message “Too slow” appeared, after which they waited the harvest delay (2 seconds total). Participants were instructed that more “too slow” messages would result in fewer apples earned. When participants harvested the patch, apples appeared on the screen for 1 second. Regardless of the reaction time, the total harvest delay was always 2 seconds long. When the participant decided to exit, the avatar character moved from the center of the screen rightwards away from the tree until it went off the screen (415 millisecond animation) then the travel task occupied the screen (7.5 seconds), after which the avatar reappeared from the left side of the screen and moved rightwards towards the tree at the center of the screen (415 millisecond animation).

### Block-wise manipulation

Patches were presented block-wise. We manipulated two factors that defined a block: effort type (cognitive and physical) and effort level (low and high). Each block type was tested twice, making 8 blocks total. The total duration of the block was fixed (4 minutes). Participants had a self-paced break between blocks. Participants were instructed that the time in a block was fixed at 4 minutes, and that they had to decide how to spend their time between harvesting and traveling (see SI Text 10 for complete instructions). The cognitive and physical variants of the task were completed separately (i.e., all cognitive effort blocks were completed in sequence, as were all physical effort blocks). Participants did not know when playing the first effort variant that there would be a second variant upcoming in the experiment. The order of cognitive and physical effort variants of the task was counterbalanced across participants. Within blocks of an effort type, each effort level was tested once during the first half and once during the second half. Given that constraint, the effort level was fully counterbalanced, resulting in eight possible block orders. Which of the block orders was used was randomly selected for each participant. Participants were explicitly instructed about which travel task they had to perform in a particular block and they could use the background color to know which travel task to perform (light blue for cognitive low effort, light orange for cognitive high effort, light purple for physical low effort, light green for physical high effort).

### Task environment

The only difference between blocks was the effort travel task, all other variables of the foraging environment were fixed (see Tab. S5). The time it took to harvest the tree (2 seconds) or travel to a new tree (8.33 seconds) were fixed, regardless of reaction times. The apple yield of the first visit to a patch was drawn from a normal distribution (*N* (15 apples, 1), maximum = 20 apples). Each following yield was the product of the previous yield and the depletion rate. The depletion rate was drawn every harvest trial from a beta distribution (*α* = 14.909, *β* = 2.033) and was on average 0.88 (the yield would not deplete below 0.5 apples).

### Travel Tasks

Participants’ bonus earnings were not influenced by their travel task performance. In the main task, we did not set a performance criterion because that would have complicated the interpretation of the foraging behavior (which would then require estimating not just effort costs but, for example, subjective efficacy estimates per participant). However, training established the expectation that participants try to be accurate by tasking participants with completing 5 mini-blocks with high accuracy. (Participants were required to repeat a mini-block if they made two errors in a row, leading the black dot to be displayed).

#### Multi-Source Interference Task

We used the Multi-Source Interference Task as the cognitive effort task. This task includes multiple types of interference effects; Stroop, Flanker, and Simon effects, and is simple to administer with a standard keyboard without the need for participants to learn novel key mappings (Bush & Shin, 2006). In this task, participants identify the oddball out of three numbers by pressing either the 1, 2, or 3 keys, the trial could either be congruent (i.e., 1 0 0, press 1), or have interference (i.e., 3 3 1, press 1). Interference trials have a competing distractor response (here, 3), for which the oddball target is flanked by the distractor and in the spatial position of the distractor (here, 1 is in the 3rd position). The MSIT trial began with 250 ms fixation cross, then the stimulus appeared for 1000 ms and participants could enter their response. After a total of 1250 ms the trial ended. Participants completed 6 trials per travel for a total of 7.5 seconds of task time. If participants made two errors in a row, they saw an attention check (black dot) for 250ms instead of the fixation cross. Participants were instructed to avoid seeing the black dot.

#### Rapid Key-pressing Task

Participants performed rapid key-pressing as part of foraging task during travel between trees (7.5 seconds, Fig. 2, see training methods in SI Text 9). In the task participants rapidly pressed the keyboard with their non-dominant pinky finger. All participants were right-handed and used their left pinky finger to press (the ‘a’ key). Each press filled a bar that spanned the horizontal extent of the screen. The horizontal bar indicated progress towards the goal number of presses. There were two conditions referred to as the “Larger number of presses” and “Smaller number of presses”. Travel time was fixed, so if participants reached the goal presses before the travel duration they waited and saw the message ‘Completed!’ on screen. If they failed to complete the goal number of presses a black dot appeared on the screen. Participants were instructed to avoid seeing the black dot. To ensure within reason that participants used their non-dominant pinky finger throughout the task, they were required to press ‘hold keys’ to occupy other fingers. The hold keys were ‘w’, ‘e’, and ‘f’ for the left hand, and ‘h’ and ‘o’ for the right hand. To minimize cognitive demands the hold keys were always displayed at the bottom of the screen during the rapid key-pressing task.

### Overview of experiment

Experiment 1 was conducted over a 90-minute session. Participants gave electronic informed consent to participate in the study. All tasks and surveys were presented using the jsPsych library for JavaScript (de Leeuw, 2015), and served with using NivTurk software (Zorowitz & Bennett, 2022) using the Flask software package for Python. Participants began the experiment with self-report surveys, followed by the foraging training, the main foraging task, and lastly a debrief survey including demographics. Based on recent theoretical work (Grahek et al., 2019) we created a battery of surveys to capture trait motivation (i.e., need for cognition, trait effortful control), and state motivation and affect (i.e., current symptoms of apathy, anhedonia, depression, anxiety, see Tab. S2).

### Participants

678 Prolific participants (18-56 years, mean=24.5 years *±* 6.7, 307 female, 365 male, and 6 prefer not to answer, race and Hispanic or Latino ethnicity reported in Tab. S6) volunteered for the study. The study was approved by the Princeton University Institutional Review Board and participants were recruited from the Prolific platform for the large online sample. Participants were compensated with $8.33 for one hour a performance bonus up to $4 (Prolific bonus mean=$3.52, standard deviation=0.78, range=$0.35 - 4). The total number of apples harvested in the Effort Foraging Task were converted into real money at the end of the experiment, with each apple being worth fractions of a cent (0.009 cents per apple). The conversion factor was set using pilot data, such that the best performing participant (earned the most apples) would make the maximum bonus. To accommodate both the physical effort task (completed with the non-dominant pinky finger) and the foraging task within standard keyboard layout, all participants were right-handed. Participants completed foraging decisions with their right hand and effort travel tasks with their left hand.

### Analysis methods

#### Hierarchical Bayesian Marginal Value Theorem Model

Constantino & Daw (2015) investigated trial-by-trial learning in the patch foraging task by predicting stay-or-exit choices using a softmax (noisy) version of a Marginal Value Theorem (Charnov, 1976) threshold rule for each stay-or-exit choice. As threshold, the model used a dynamic reward rate estimate given by a running average over obtained rewards and experienced delays. They showed that this model outperformed other candidate learning rules, notably temporal difference learning.

For the present study, because we are investigating individual differences in effort costs at the condition level, we simplified that model to a factorial one in which the MVT threshold is instead taken as fixed per-condition, determined by the overall rewards and delays in each condition and a per-condition effort-cost parameter. Thus, the model omits trial-by-trial learning of the threshold, and instead formally absorbs any such variation into the softmax choice stochasticity. We believe this simplification is warranted because the condition-wise effort costs of interest aggregate over per-trial threshold variability, and because we encouraged asymptotic behavior through extensive pre-training and using a stable foraging environment throughout. Also, we have found that learning effects are more easily estimated in a different class of foraging tasks, prey-selection tasks (Krebs et al., 2010; Blanchard & Hayden, 2014; Garrett & Daw, 2020; Toro-Serey et al., 2021), because trial-to-trial prey encounters are independent, whereas the decaying reward dynamics in patch foraging tasks correlate offers across trials.

In these respects, this model is intended as descriptive rather than as a process model. Indeed, our approach of solving for the effort costs that rationalize asymptotic behavior is also compatible with alternative assumptions about the way the decision variables are computed (e.g., prospectively, as an intertemporal choice between anticipated future rewards minus costs from harvesting vs. traveling).

First, we computed known reward rate values of the foraging environment per effort condition per participant: total rewards harvested (Σ*r*), number of harvest periods (*T* =block duration/harvest time), and total times travelled (see foraging environment parameters in Tab. S5). Then, we solved for the unknown component of average reward rate; the cost of travel (*c*). We estimated the cost of the high effort task (*c_high effort_*) for an individual by predicting harvest versus exit decisions using a hierarchical Bayesian logistic model (equations 2 to 5). For each foraging trial, model compares the expected reward on the next harvest (*R_e_*, defined as the average of the previous harvest and the product of the previous harvest with the mean depletion rate (0.88)) against the overall average reward rate for a block type (*ρ*), using a softmax function (with inverse temperature parameter, *β*) to make a choice (harvest or exit). The cost of travel in high effort blocks (*c_high effort_*) was expressed as the marginal increase in cost of travel (*c_low effort_* + *c_high effort_*) from low to high effort. Defining this cost as a difference measure controls for any additional biases individual participants may have which are common to both conditions (i.e., consistently high exit thresholds for some participants and low thresholds for others). We used (*c_high effort_*) as the dependent measure of the effort cost for an individual.

For each effort level (low and high) and effort type (cognitive and physical) we predicted choices to stay or exit a patch:

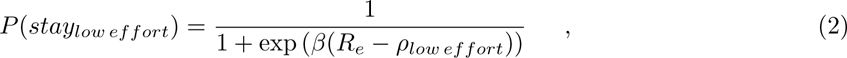

where,

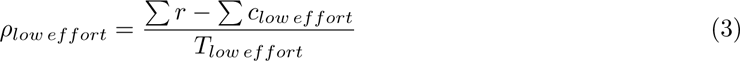

and,

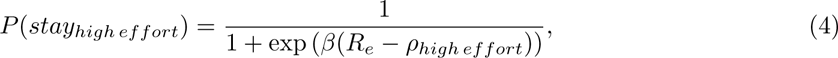

where,

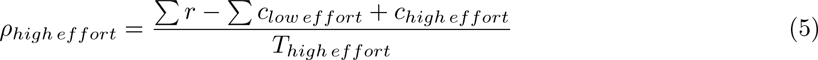

There were five parameters in the model, the inverse temperature (*β*, which controls the noise of the softmax choice function, with lower values indicating more noisy effects of rewards and thresholds on choices), the cognitive low (*c_cog low effort_*) and high effort costs (*c_cog high effort_*), and the physical low (*c_phys low effort_* and high effort costs (*c_phys high effort_*). The model included a full covariance matrix of the parameters (5-by-5 matrix) which consists of a correlation matrix and a scale (standard deviation) matrix. Parameters were drawn from a multi-variate Gaussian distribution. We used the covariance matrix to estimate the correlation between individual differences in high cognitive and physical effort costs.

The prior distributions were *c_low effort_ ∼ N* (0, 40), *c_high effort_ ∼ N* (0, 30), *β ∼ N* (0, 0.5). The prior on the correlation matrix was unbiased as to the presence or absence of a correlation (LKJ Correlation Distribution prior=1, (Lewandowski et al., 2009)). Individual participant parameters and their group-level distributions were estimated using Markov Chain Monte Carlo sampling, implemented in Stan with the CmdStanR package (4,000 samples, 2,000 warm-up samples, across 4 chains, Stan Development Team, Stan, 2021). We also simulated the MVT model to estimate the best exit threshold with respect to reward and time given the foraging environment parameters (Tab. S7, SI Text 6).

#### Canonical correlation analysis

To leverage the strength of our data in having many detailed individual differences measures of theoretically related constructs we used Canonical Correlation analysis to perform a many to many correlation (we used the cc function from the CCA package in the R language, Gonález & Déjean, 2021). The task measures included were cognitive effort cost, interference and congruent trial error rate (transformed as log(2-correct)), physical effort cost, and overall exit threshold (estimated in log apples over all low effort blocks by participant using linear mixed-effects regression, discussed in SI Text 1). The self-report measures were a combination of trait and symptom state measures of motivation for cognitive and physical effort (SI Text 4, Tab. S2). The trait self-reports were the need for cognition (Cacioppo et al., 1984), behavioral inhibition and behavioral activation (Carver & White, 1994), effortful control (Adult-Temperament Questionnaire Evans & Rothbart, 2007). The symptom self-reports were cognitive function (PROMIS, Cella et al., 2007), apathy (Apathy Motivation Index Ang et al., 2017), anhedonia (Snaith Hamilton Pleasure Scale, Snaith et al., 1995), physical fatigue (PROMIS Fatigue, Cella et al., 2007), general self-efficacy (PROMIS, Cella et al., 2007), anxiety (Generalized Anxiety Disorder-7, Spitzer et al., 2006), and depression (Patient Health Questionnaire-9 Kroenke et al., 2001). For any measures with subscales, self-report scores were the combined overall averages.

#### Exclusion criteria

Participants completed the study on their own outside of the laboratory. To ensure data quality we used task behavior to constrain our sample to participants who completed the experiment in earnest (SI Text 7, Tab. S8 shows the Experiment 1 exclusion criteria, the quantitative cutoffs, and the number of outlier participants). The exclusion criteria were not completing the experiment, missing the response deadline on many harvesting trials, poor cognitive or physical travel task performance, too few exit trials in a condition, and very large changes in exit threshold from low to high cognitive and physical effort conditions. Participants with very large shifts in thresholds produced strong outliers in our Marginal Value Theorem model, many of whom had very few exit trials in one condition.

### Code and Data availability

The Effort Foraging Task code, the analysis code, and data are openly available at the Open Science Framework repository ‘Effort Foraging Task’, https://osf.io/a4r2e/.

## Acknowledgements

Thanks to Sara Constantino for providing base task code for the in-laboratory experiments and Evan Russek for providing base task code for the remote experiments. Thanks to Allison Burton, Temitope Oshinowo, and Connor Lawhead for collecting the in laboratory data. Thanks to Elizabeth Tong, Temitope Oshinowo, and Jeremy Lee for adapting and developing online task code. Thanks to Kevin Lloyd for providing the base code for the optimal threshold simulations. Thanks to Sebastien Bouret for helpful comments regarding the relationship between cognitive and physical effort costs. Thanks Yoel Sanchez Araujo, Sam Zorowitz, Ben Singer and Dave Turner for assistance with cluster computing for model fitting. Thanks to Lindsay Hunter for feedback on the manuscript. This work was supported by the John Templeton Foundation, the National Science Foundation Graduate Research Fellowship Program, and the National Institutes of Health Grant T32MH065214. The opinions expressed in this publication are those of the authors and do not necessarily reflect the views of the John Templeton Foundation.

## Supporting Information Text

### 1. Overall threshold results

We computed the group mean overall exit thresholds separately in Experiments 1 and 2 using mixed effect linear regression using only an intercept term. We compared these observed group overall thresholds to best thresholds from simulations (see group-level means by condition in Fig. S2). We found that the mean exit threshold across all conditions in Experiment 1 was 6.30 apples (SE = 0.11, df = 615.95, t = 56.09, p < 0.001). The group average was close to the best threshold identified by simulation (6.78 apples), however individuals varied widely. The group-level mean overall threshold was 4.02 apples in Experiment 2 (SE = 0.22, df = 80.22, t = 18.47, p < 0.001) which was close to the best policy in simulation of 4.65 apples. Using linear regression we estimated the mean exit threshold across all conditions (“overall threshold”) per participant and included these estimates as an additional dimension of individual differences in the task (when testing the relationship between task behavior and surveys). The overall exit threshold may be a relevant individual difference in representations of subjective reward rate (Fig. S2). A benefit of this task is that it can simultaneously measure exit thresholds and effort costs. Striatal dopamine is hypothesized to represent average reward rate in foraging settings (Constantino et al., 2017; Le Heron et al., 2020). Consistent with this exit thresholds are lower (more over-harvesting) in individuals with Parkinson’s (Constantino et al., 2017; Le Heron et al., 2018), when participants are chronically or acutely stressed (Lenow et al., 2017), and in individuals with opioid dependence (Raio et al., 2022). Critically, high cognitive and physical effort costs are derived from a shift in exit threshold, this partials out the variance having to do with overall threshold.

**Fig. S1.**
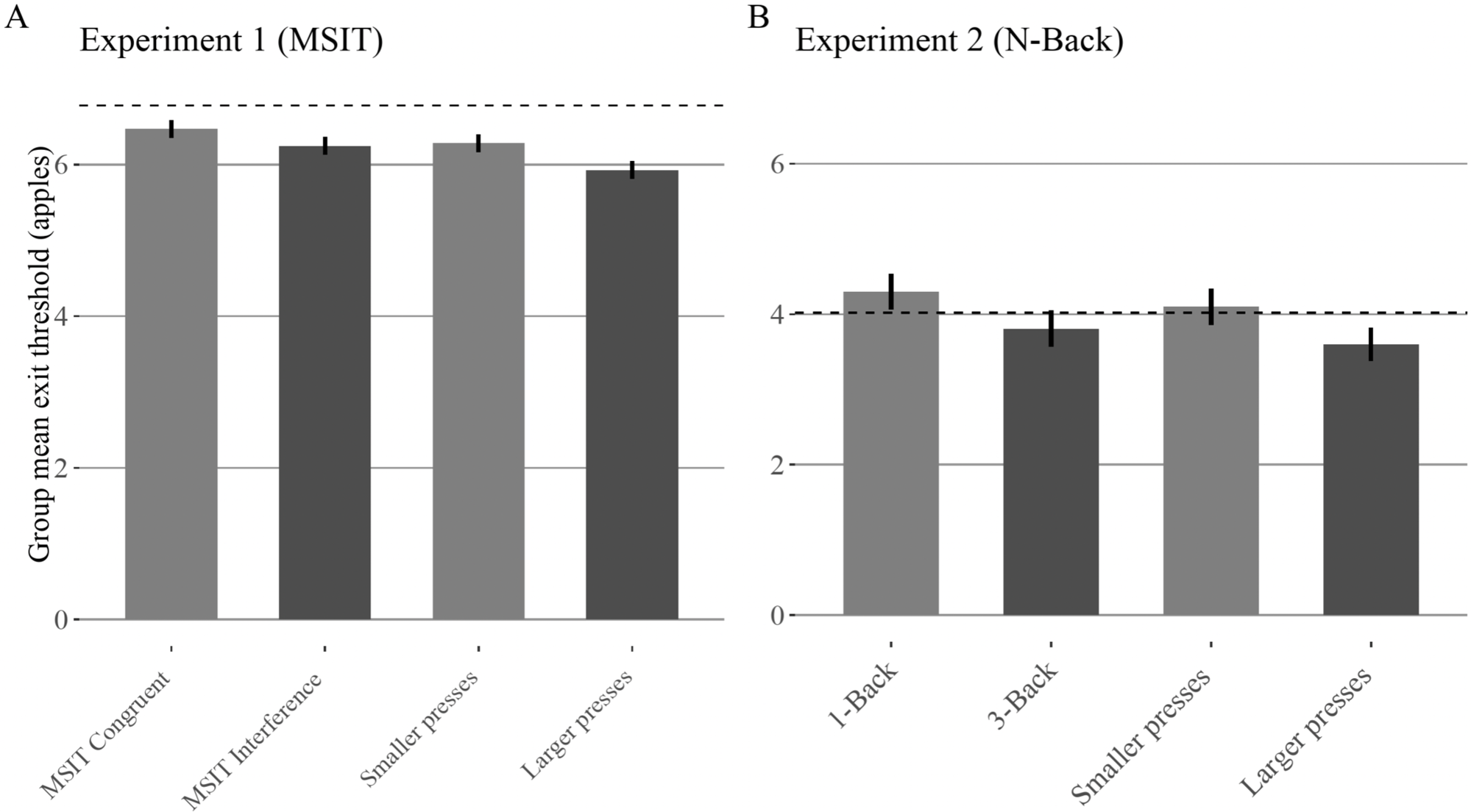
Group-level exit thresholds Experiment s 1 and 2. A: Experiment 1, B: Experiment 2. x-axis: Foraging conditions indicated by the travel task required, y-axis: Group-level mean exit threshold (apples), error bars indicate SEM. Effort level indicated by bar color (light gray = low effort, dark gray = high effort). Group average thresholds near best threshold from simulation (best threshold respect to reward rate indicated by dotted line, 6.78 apples in Experiment 1 (MSIT), 4.65 apples in Experiment 2 (N-Back)).

**Fig. S2.**
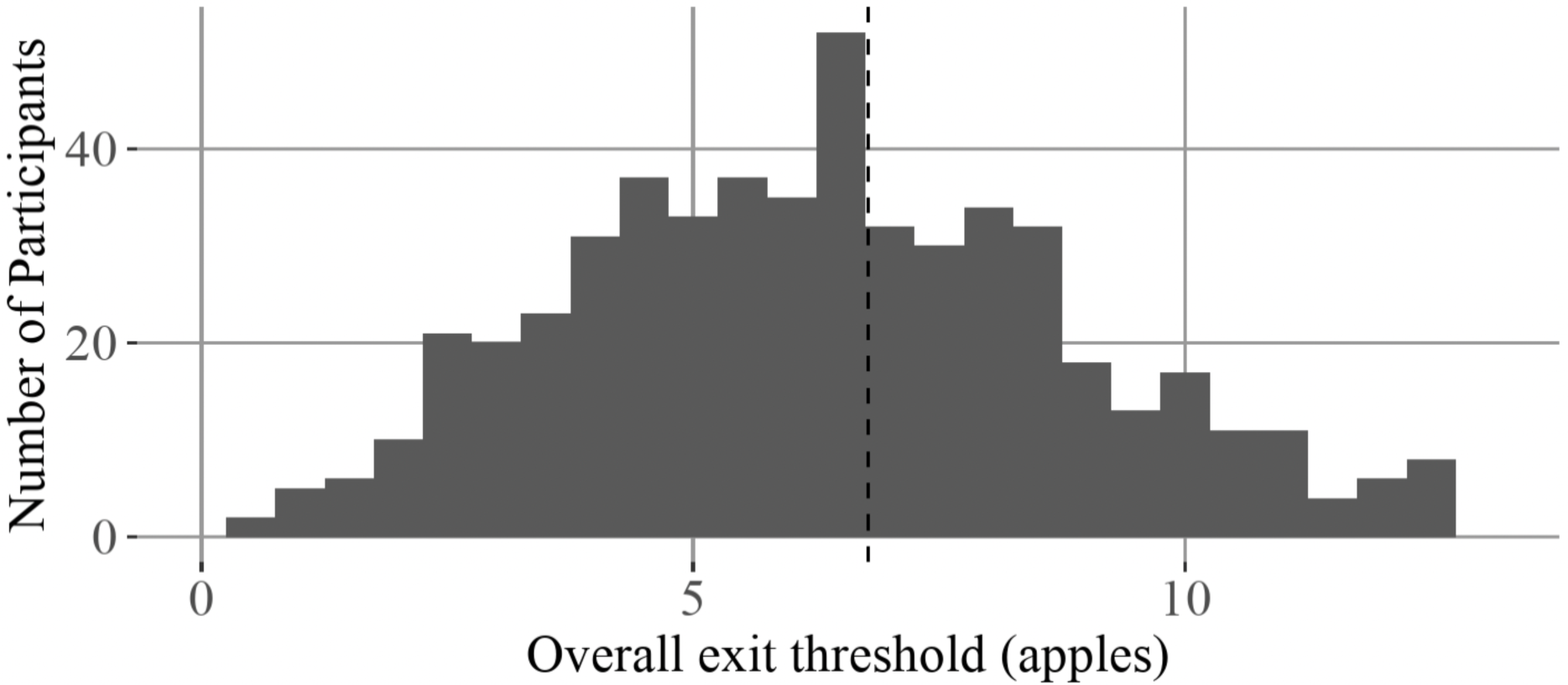
Individual differences in overall threshold. Experiment 1 histogram of individual differences in mean foraging exit threshold estimated using mixed-effects regression Experiment 1 (MSIT). Some individuals over-harvest (exit threshold below best threshold, dotted line, 6.78 apples) while others under-harvest (exit threshold above best threshold).

**Table S1.** Group level posterior distributions. The Experiment 1 (MSIT) group level average high cognitive effort cost was 7.5 apples. The group level average high physical effort cost was 13.5 apples. For each parameter (column 1) the table shows the mean of the group-level posterior distribution (column 2) and the 95% highest density interval (lower bound, column 3; upper bound, column 4).

| Parameter | Mean | Lower bound | Upper bound |
| --- | --- | --- | --- |
| Cognitive Effort Cost | 7.547 | 5.472 | 9.618 |
| Cognitive Low Effort Travel Cost | 94.646 | 90.704 | 98.746 |
| Physical Effort Cost | 13.527 | 10.903 | 15.883 |
| Physical Low Effort Travel Cost | 99.781 | 95.399 | 103.838 |
| Inverse Temperature | 0.260 | 0.239 | 0.283 |

### 2. Split-half reliability

We computed the split-half reliability of the change in exit threshold measures in Experiment 1 using hierarchical regression modeling following Rouder & Haaf (2019). We took all the exit trials in a single block and split them in half, we adapted the mixed-effects regression model to fit the change in exit threshold separately for the first and second halves, then we examined the random effects correlation of the first and second half change exit threshold. The resulting reliability was r=0.85 for cognitive effort and 0.82 for physical effort.

### 3. Cognitive and physical effort cost relationship excluding negative effort cost participants

The consistent observation of participants showing negative effort costs is interesting and opens up a research agenda on effort seeking using this task. However the mechanisms underlying effort seeking remain to be understood. Did negative effort costs reflect value added by effort, boredom aversion, or another unmeasured individual difference? We confirmed that the Experiment 1 finding of a positive correlation between cognitive and physical effort costs was maintained within the effort-avoiding group of participants, by rerunning the MVT model excluding participants with negative cognitive (N=78) or physical (N=67) effort costs (i.e., effort-seekers). This left 428 effort-avoiding (i.e., positive effort cost) participants. The positive correlation between cognitive and physical effort costs was robust to this exclusion of participants exhibiting negative effort costs (Fig. S3, 95% highest density interval is −0.021 to 0.851, mean=0.406, and 94.5% of samples>0).

**Fig. S3.**
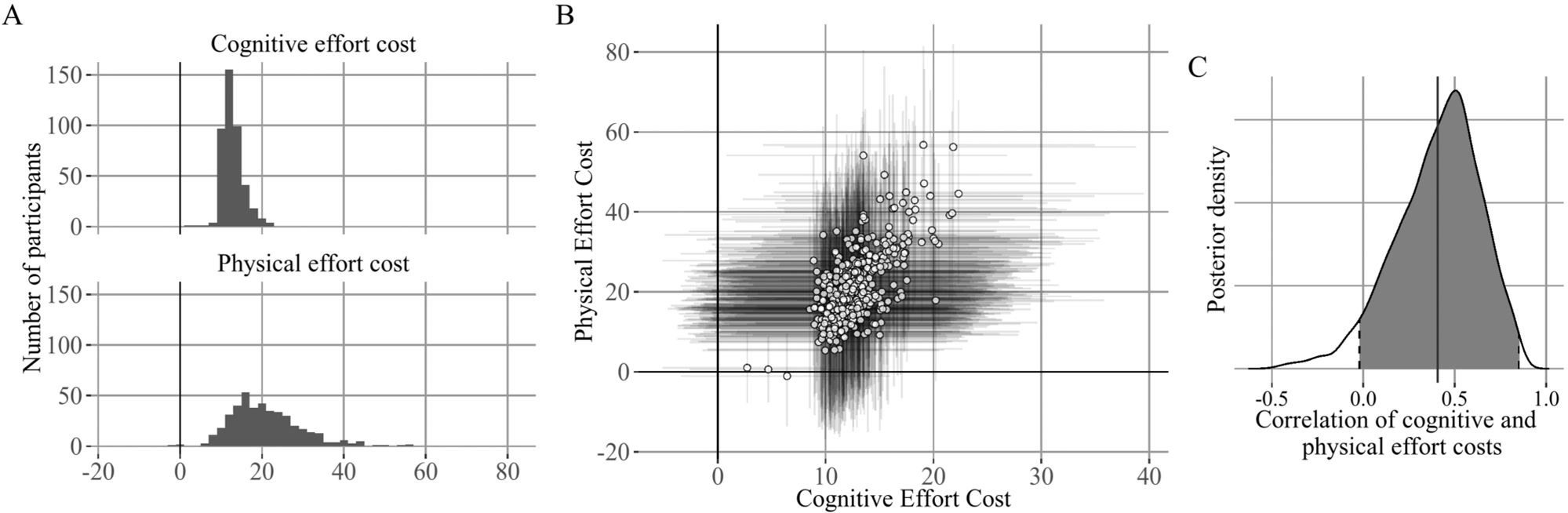
Cognitive and physical effort cost relationship excluding negative effort cost participants. Experiment 1 correlation between individual differences in cognitive and physical effort costs. A: Individual differences in the high effort travel costs (expressed as the additional cost of the high relative to the low effort condition). Paralleling the pattern of exit thresholds most participants experienced the high effort conditions as effortful (positive cost), whereas some participants were insensitive to the effort manipulation (cost near zero) and others were effort seeking (negative cost). B: x-axis: Individual differences in cognitive effort costs, y-axis: Individual differences in physical effort costs. Error bars indicate 80% HDI. C: posterior distribution of correlation between high effort cost for cognitive and physical effort. Cognitive and physical effort costs are positively correlated.

**Fig. S4.**
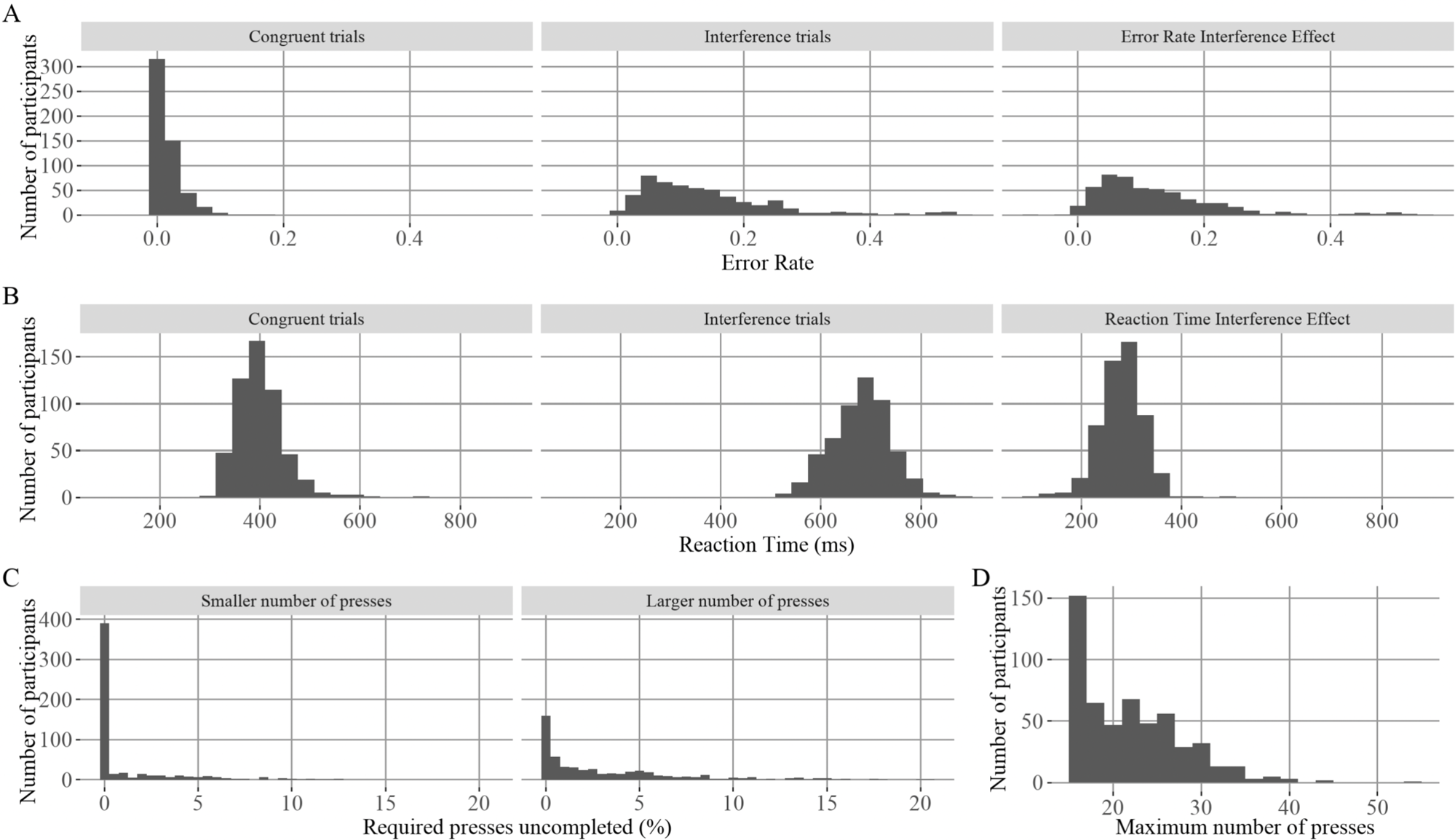
Individual differences in travel task performance. Histogram of individual differences in travel task performance for Experiment 1 participants (after exclusions). A: Error rate (computed as log(2-correct)), B: Reaction time. Column 1: congruent trials, column 2: interference trials, column 3: interference effect (interference minus congruent). C: Rapid key pressing performance. Column 1: uncompleted smaller number of presses (% of required presses), column 2: uncompleted large number of presses, column 3: maximum number of presses determined in calibration phase (15 presses was set minimum).

**Fig. S5.**
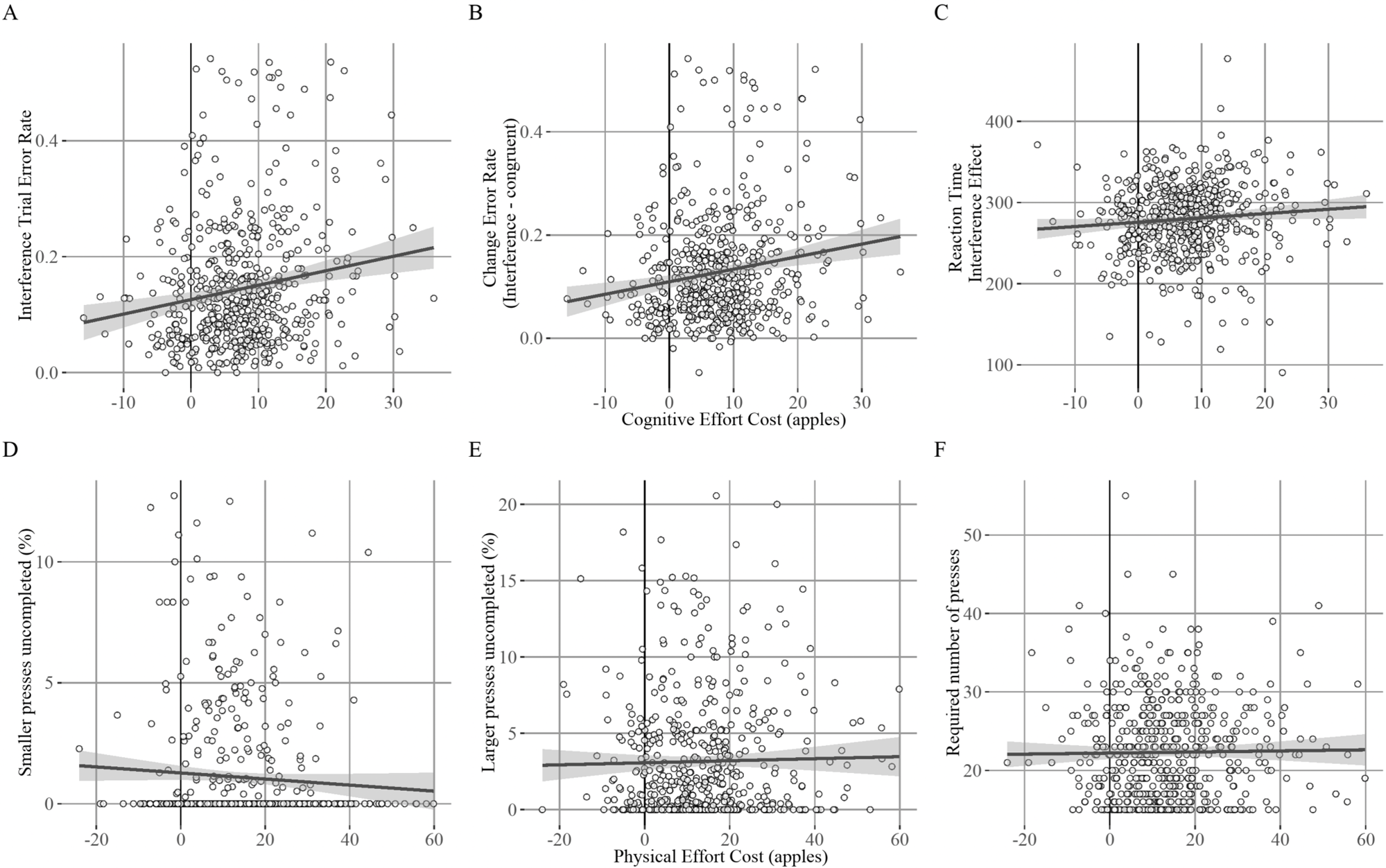
Relationship between travel task performance and effort costs. Experiment 1 cognitive effort cost positively related to error rate measures but not reaction time interference effect. A: Interference trial error rate (log transformed), B: change in error rate (Interference - Congruent), C: change in reaction time (Interference - Congruent). Physical effort cost not related to rapid keypressing performance. D: percent of smaller presses uncompleted. E: percent of larger presses uncompleted. F: required number of presses determined in calibration phase.

### 4. Self-report surveys

In Experiment 1 we collected a number of measures of current experience with psychiatric symptoms. The Apathy Motivation Index (Ang et al., 2017) which measures apathy and motivation in the behavioral, social, and emotional domains and was designed to be suitable for use in the general population. The Snaith–Hamilton Pleasure Scale (SHAPS, Snaith et al., 1995) measures anhedonia by asking about responses to common domains of pleasure. We administered the Patient Health Questionnaire-9 (PHQ-9, Kroenke et al., 2001), a common measure of depression symptoms, and the Generalized Anxiety Disorder-7, a common measure of anxiety symptoms (Spitzer et al., 2006). We administered four scales from the Patient-Reported Outcomes Measurement Information System (PROMIS, Cella et al., 2007); the Cognitive Function Short Form 4a, the Cognitive Function Abilities Short Form 4a, Fatigue Short Form 4a, and General Self-Efficacy. Given our particular interest in cognitive function symptoms such as slowed thinking and reduced concentration we used the PROMIS Cognitive Function Short Form 4a which measures subjective cognitive functioning. For this subscale higher scores indicate fewer complaints about recent cognitive function. We also used the complimentary PROMIS Cognitive Function Abilities Short Form 4a, for which higher scores indicate better subjective cognitive function. The combination of these subscales provides a total PROMIS Cognitive Function score for which higher values indicate better recent cognitive function. Lastly, we used the PROMIS Fatigue Short Form 4a to measure physical fatigue symptoms which we hypothesized would be correlated with greater physical effort costs.

In addition to symptom state measures, we also collected several trait measures. To capture self-reported cognitive control capacity, we used the Adult Temperament Questionnaire - Effortful Control scale (Evans & Rothbart, 2007) which had been related to depression in a previous study (Marchetti et al., 2018). We also used the Need for Cognition scale (Cacioppo & Petty, 1982; Cacioppo et al., 1984) which measures the extent to which individuals are prone towards engage in cognitively effortful activities. We predicted a negative relationship between cognitive effort cost and need for cognition given previous reports of such a relationship (Westbrook et al., 2013). We collected the Behavioral Inhibition, Behavioral Activation Scales (BIS/BAS, Carver & White, 1994) which measure an individuals’ sensitivity to the behavioral approach and behavioral avoidance system (abbreviated form Pagliaccio et al., 2016). This measure was useful for several purposes. Firstly, we used it as a measure of reward sensitivity, in line with evidence linking Behavioral Activation to striatal activation in anticipation of rewards (Costumero et al., 2016). Research also shows Behavioral Activation scores tend to be lower and Behavioral Inhibition scores tend to be higher in more depressed participants (Kasch et al., 2002; McFarland et al., 2006; Pinto-Meza et al., 2006; Alloy et al., 2008; Quilty et al., 2014). The theoretical work by Grahek and colleagues (2019; Grahek et al., 2018) also identified self-efficacy as a potentially relevant factor in cognitive control decision making in depression. By this account more depressed participants may be less likely to predict that exerting effort will lead to a rewarding outcome. We used the PROMIS General Self-Efficacy scale, in which participants respond how confident they are for items such as ‘I can manage to solve difficult problems if I try hard enough’. We predicted participants with higher self-efficacy would have a stronger belief that effort will result in reward, potentially leading to a greater propensity to exert effort. Though, ideally this factor would be assessed with a cognitive task that manipulates efficacy (see Frömer et al., 2021).

#### A. Self-report attention checks

Infrequent attention check items were embedded in the self-report surveys in Experiment 1 to ensure participants were reading the items (following Zorowitz et al., 2021). 28 participants were excluded from self-report analyses (i.e., CCA) because they failed either of the attention check items embedded in the Apathy Motivation Index, and Generalized Anxiety Disorder-7 self-reports (Fig. S6). We did not use the items embedded in PROMIS-Cognitive Function or the Patient Health Questionnaire-9 self-reports because response patterns indicated ambiguity in the questions.

**Fig. S6.**
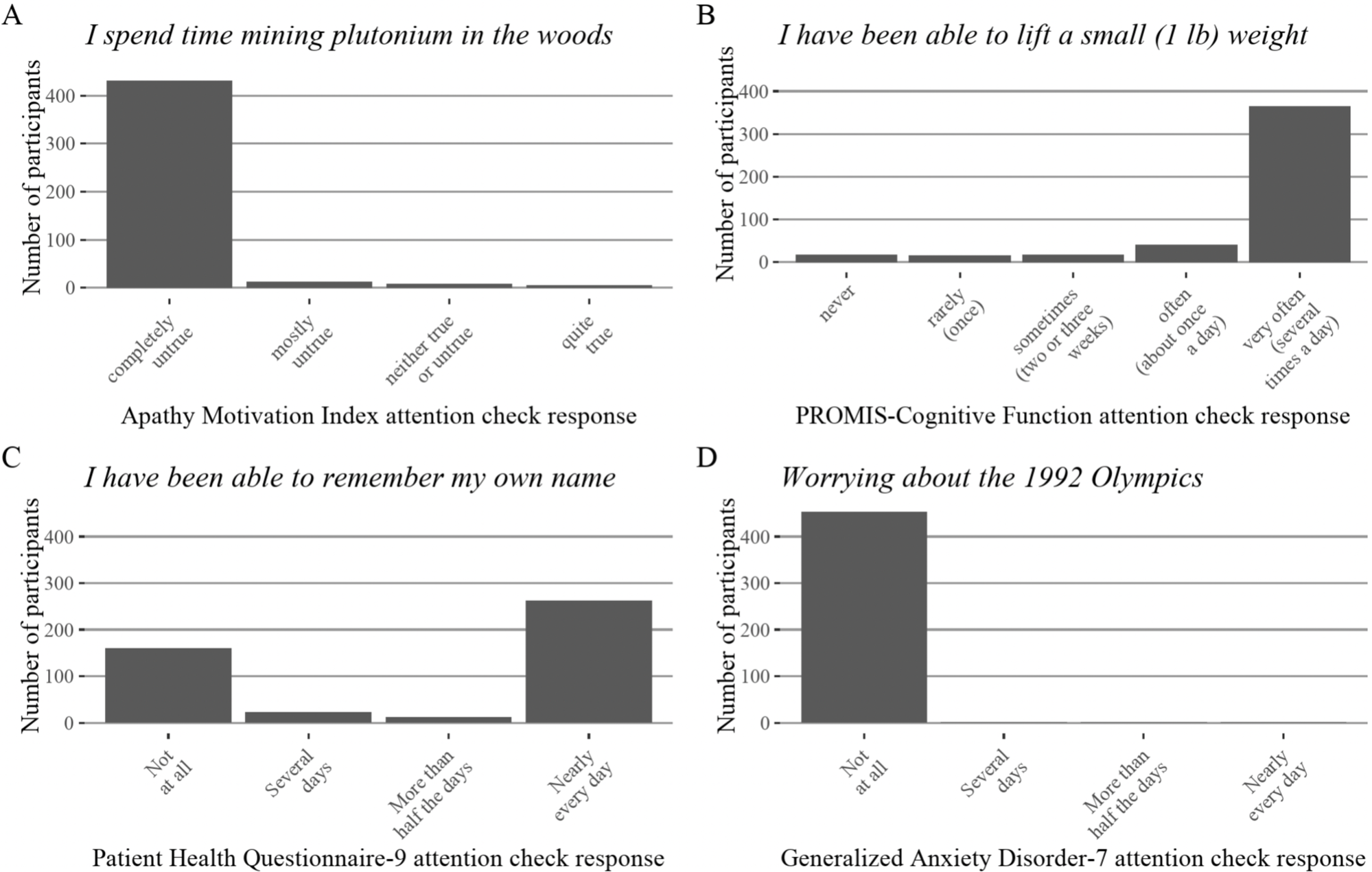
Attention check items results. A: Apathy Motivation Index, “I spend time mining plutonium in the woods”, correct answer “completely untrue”. B: PROMIS-Cognitive Function, “I have been able to lift a small (1 lb) weight, correct answer “very often (several times a day)”. C: Patient Health Questionnaire-9, “I have been able to remember my own name”, correct answer “Nearly every day”. D: Generalized Anxiety Disorder-7, “Worrying about the 1992 Olympics”, correct answer “Not at all”. Experiment 1 participants were excluded if they were inattentive on the Apathy Motivation Index, and Generalized Anxiety Disorder-7 items.

**Table S2.** Self-report survey battery. Column 1: scale name; column 2: number of items; column 3; construct measured and variable name used in the canonical correlation analysis; column 4; timescale of survey instructions, ‘trait’ indicates surveys about behaviors characteristic of the individual. Experiment 1 self-reports were completed in the order listed in this table. Scales asking about similar timescales were grouped together.

| Scale name | N items | Construct measured | Time scale |
| --- | --- | --- | --- |
| Patient Health Questionnaire-9 | 9 | Depression | last 2 weeks |
| Generalized Anxiety Disorder-7 | 7 | Anxiety | last 2 weeks |
| Apathy Motivation Index | 14 | Apathy | last 2 weeks |
| Snaith-Hamilton Pleasure Scale | 13 | Anhedonia | last few days |
| PROMIS Cognitive Function Short Form 4a | 4 | Cognitive Function | last 7 days |
| PROMIS Cognitive Function Abilities Short Form 4a | 4 | Cognitive Function | last 7 days |
| PROMIS Fatigue Short Form 4a | 4 | Fatigue | last 7 days |
| PROMIS General Self-Efficacy | 4 | Self-efficacy | last 7 days |
| Need for Cognition | 18 | Need for Cognition | trait |
| Adult Temperament Questionnaire-Effortful Control | 19 | Effortful Control | trait |
| Behavioral Inhibition/Activation Scales - Abbreviated | 12 | Behavioral Inhibition/Behavioral Activation | trait |
| **Total** | 108 |  |  |

**Fig. S7.**
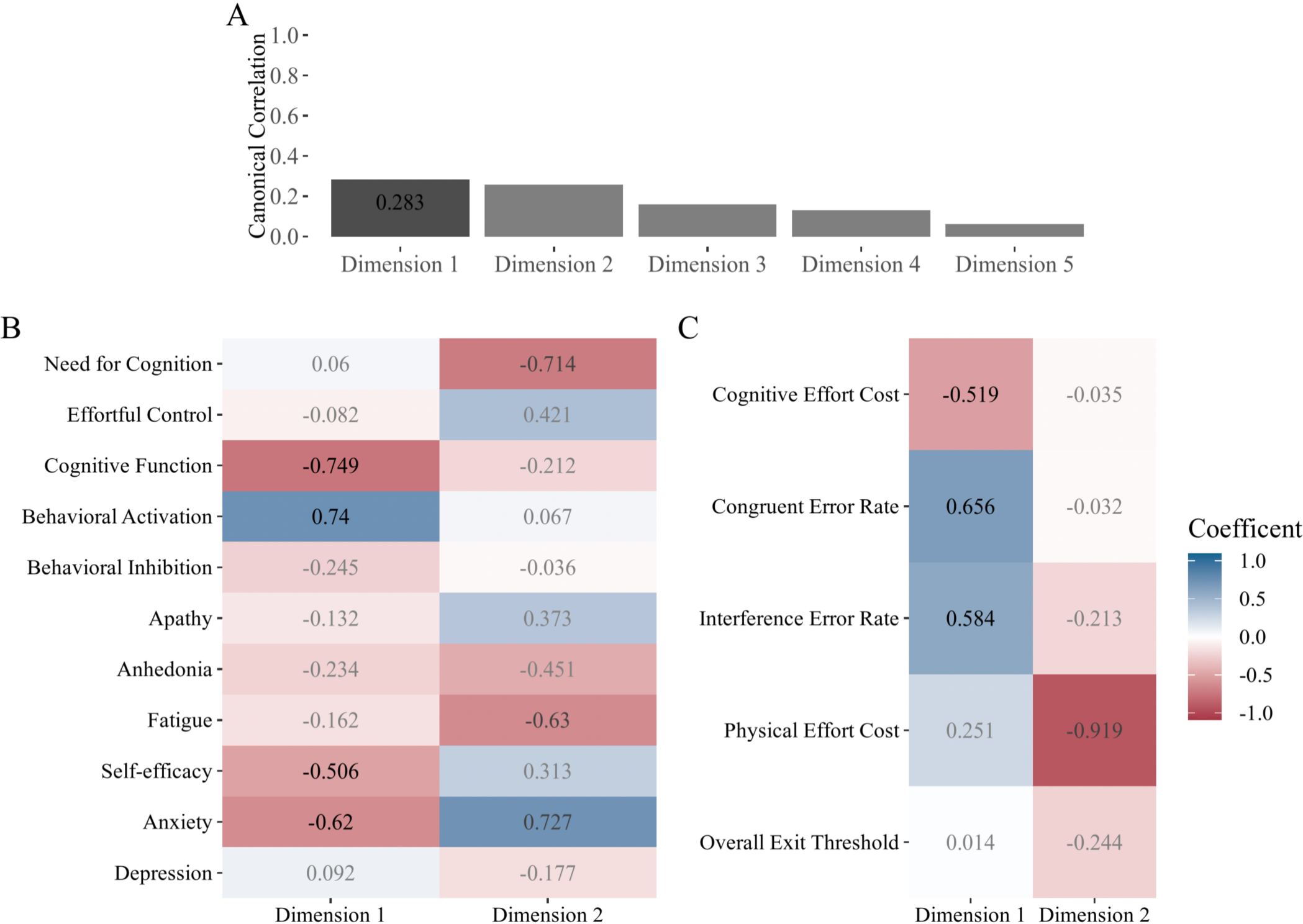
Experiment 1 full canonical correlation results. A: canonical correlations by dimension. B: dimension coefficients for task parameters (X coefficients). C: dimension coefficients for self-reports (Y coefficients). Coefficients with absolute value larger than 0.5 from significant dimensions are shown in black for the significant dimension and dark gray for non-significant dimension, while coefficients below threshold are shown in light gray. Only the first two dimensions are displayed.

### 5. Validation Experiments

#### A. Experiment 2 (N-Back)

We developed two cognitive effort variants of the effort foraging task. In both versions we used a cognitive and a physical effort manipulation. Experiment 2 (N-Back) was developed in an undergraduate population. Experiment 1 (MSIT) was an abbreviated version developed for a large-scale online study. This tested the generalizability of the task in terms of population as well as in the type of cognitive effort (working memory versus inhibition).

##### A.1. Participants

116 Undergraduate students volunteered for a 2.5-hour self-guided remote experiment (Experiment 2 (N-Back), 18-27 years, mean=20 years *±* 1.5, 70 female, 42 male, 4 prefer not to answer). The study was approved by the Princeton University Institutional Review Board and participants were recruited from a pool maintained by the Princeton Psychology Department. Undergraduate students were compensated with 2.5 psychology course credit hours and a performance bonus up to $10 in the form of an Amazon gift card (bonus mean=$7.68, SD=0.61, range=$4.41 - 8.35). The conversion of apples to money was 0.11 cents per apple. Participants were excluded from analysis based on their behavior in the task (Tab. S3).

##### A.2. N-Back working memory task

The N-Back task was performed as part of foraging task during travel between trees. In the N-Back task letters are displayed on screen in a sequence. Participants judged whether the stimulus that is currently on the screen matches the stimulus they saw a number of screens back (N-Back). On every trial, participants responded whether the letter was a match (“s” key) or non-match (“d” key) to the letter on the previous screen (1-Back case) or three screens before (3-Back case). A trial began with a fixation cross (for 250 milliseconds) followed by the letter on screen (for 500 milliseconds) followed by a blank screen (for 950 milliseconds, total trial duration = 1.7 seconds). During the travel period, 10 letters were presented, of which, 2 or 3 were targets (letter matches letter N-Back) and 2 or 3 were lures (matches current letter but not in position N-Back). The number of targets and lures were selected randomly each time an N-Back stimulus sequence was generated. We only used consonants to prevent participants from using mnemonics (letters were: ‘B’, ‘C’, ‘D’, ‘F’, ‘G’, ‘H’, ‘J’, ‘K’, ‘M’, ‘N’, ‘P’, ‘Q’, ‘R’, ‘S’, ‘T’, ‘V’, ‘W’, ‘X’, ‘Y’, ‘Z’), and half of the letters were presented in upper case and the other half lower case to prevent participants using iconic memory (Cohen et al., 1994).

##### A.3. N-Back working memory task training

We trained the N-Back task extensively to try to bring participants to highest possible levels of performance and minimize automaticity differences (in which some participants would have more experience with the N-Back or similar tasks, making the task less effortful for them compared to someone with little experience). Participants had to reach a performance criterion to move on from training. After being instructed on the task participants began practice for one of the effort levels (counterbalanced). First, they completed two extended blocks (50 trials with a self-paced break up to 45 seconds between) with feedback about error type (types of feedback: “non-match”, “missed match”, “no response”, displayed in red font for 800 ms after the trial). Then they performed one extended block without any feedback (50 trials).

We tasked participants with completing a set number of mini-blocks with high accuracy to begin the foraging task. We did so to establish the expectation that participants had to exert effort when they chose to travel while foraging. A mini-block was classified as successful when the participant saw no error feedback (large black dot), after which they were told they were moving on to the next mini-block. The error feedback was displayed when participants made two consecutive errors (including omission errors). If they did see one or more error feedback symbols, they had to repeat that mini-block. They had to successfully complete 8 mini-blocks of the 1-Back task, and 12 mini-blocks of the 3-Back task. This training also ensured that participants could adequately perform the task. Participants had self-paced breaks in between mini-blocks (up to 60 seconds).

##### A.4. No relationship between cognitive and physical effort cost in Experiment 2

In the larger online sample, we found a significant positive correlation between cognitive and physical effort costs (Experiment 1 (MSIT), N=537, correlation=0.55). In the smaller Experiment 2 (N-Back) (N=81) sample we did not find conclusive evidence for or against the correlation, as the highest density interval (HDI) was very wide (Fig. S8, mean correlation=0.048, 95% HDI=-0.369 - 0.462). One explanation is that we were underpowered to detect a correlation similar in size to that seen in Experiment 1. Encouragingly, the HDI for the Experiment 2 (N-Back) model overlapped with the HDI in Experiment 1 (MSIT). Another difference between experiments is that Experiment 2 is an undergraduate student population, while Experiment 1 might more closely reflect the general population. We speculated that, with respect to our experimental question ‘are cognitive and physical effort cost correlated?’, there may be a selection bias due to conditioning on a collider in Experiment 2. The collider would be admission to university (a classic example Hernán & Monge, 2023), which may select for students specializing in either academics (cognitive effort) and not athletics (physical effort) or vice versa. This negative correlation induced in this population between academic and athletic ability may tend to cancel out the positive correlation demonstrated in Experiment 1. This could be tested in a larger population of undergraduates sufficient to confidently detect the presence or absence of a correlation.

##### A.5. Explicit awareness of effort avoidance

In Experiment 2 (N-Back), we asked participants if, and how, the required travel task changed their decision to travel to a new tree. For the cognitive (N-Back) variant, of those who completed the debrief survey (N=113), 36% of participants (N=41) reported changing their behavior based on the travel task (saying in their own words that they avoided the high cognitive effort [3-Back] task and stayed longer at a tree), whereas 64% of participants (N=72) explicitly stated that the travel task did not change their decisions. This supports the idea that the task is an indirect measure for the majority of participants (i.e., participants whose behavior was influenced by the travel task were not aware of doing so).

##### A.6. Effort ratings

We asked participants to rate how effortful each of the travel tasks was. For Experiment 2 (N-Back), 9 participants reported no change in effort rating between high and low cognitive effort, 2 participants reported the low effort task as more effortful, and the remaining 102 participants rated the high effort task as more effortful. For Experiment 2 (N-Back), 17 participants reported no change in effort rating between high and low physical effort, 1 participant found the low effort task (smaller presses) to be more effortful, and the remaining 95 participants reported the high effort task (larger presses) was more effortful. On average, effort ratings were higher for the 3-Back than the 1-Back task (paired two-tailed t-test, mean of the differences=1.99, t=19, df=112, p<0.001). Participants also found the larger button presses condition to be more effortful than the smaller presses on average (mean of the differences=1.48, t=15.42, df=112, p<0.001).

**Table S3.** Experiment 2 (N-Back) Exclusion Methods. Column 1: basis of exclusion, column 2: numbers of outlier participants for Experiment 2 (N-Back), column 3: exclusion cutoff value. Participants could be excluded on multiple grounds, therefore the number of outlier participants listed are do not reflect the total number of excluded participants (shown in the bottom two rows).

| Exclusion criteria | N outlier participants | Cutoff | Mean, SD |
| --- | --- | --- | --- |
| Did not complete experiment | 9 |  |  |
| Foraging trial response deadline missed (> 10% trials) | 7 | 10 | (4.933, 13.893) |
| 1-Back accuracy ($D'$, < 2SD) | 5 | 1.723 | 3.070, 0.675 |
| 3-Back accuracy ($D'$, < 2SD) | 5 | 0.665 | 2.260, 0.796 |
| Smaller number of presses uncompleted (% , >2SD) | 9 | 2.827 | 0.4428, 1.192 |
| Larger number of presses uncompleted (% , >2SD) | 5 | 8.213 | 1.777, 3.218 |
| Number of exit trials per condition (<2SD) | 12 | 8.268 | 17.767, 4.749 |
| Change exit threshold cognitive (apples, <2SD \| >2SD) | 5 | (<-3.1145 \| >1.967) | -0.574, 1.270 |
| Change exit threshold physical (apples, <2SD \| >2SD) | 6 | <-2.964 \| >1.725 | -0.619, 1.172 |
| Initial Sample Size | 116 |  |  |
| Final Sample Size | 81 |  |  |

**Table S4.** Experiment 2 (N-Back) Parameter posterior distribution values. Table includes the mean of the group-level posterior distribution and the upper and lower bounds (95% HDI).

| **Parameter** | **Mean** | **Lower bound** | **Upper bound** |
| --- | --- | --- | --- |
| Inverse Temperature | 0.425 | 0.383 | 0.469 |
| Cognitive Low Effort Travel Cost | 114.909 | 105.088 | 125.274 |
| Cognitive Effort Cost | 16.776 | 10.356 | 23.441 |
| Physical Low Effort Travel Cost | 124.295 | 112.371 | 136.350 |
| Physical Effort Cost | 15.943 | 8.915 | 23.007 |

**Fig. S8.**
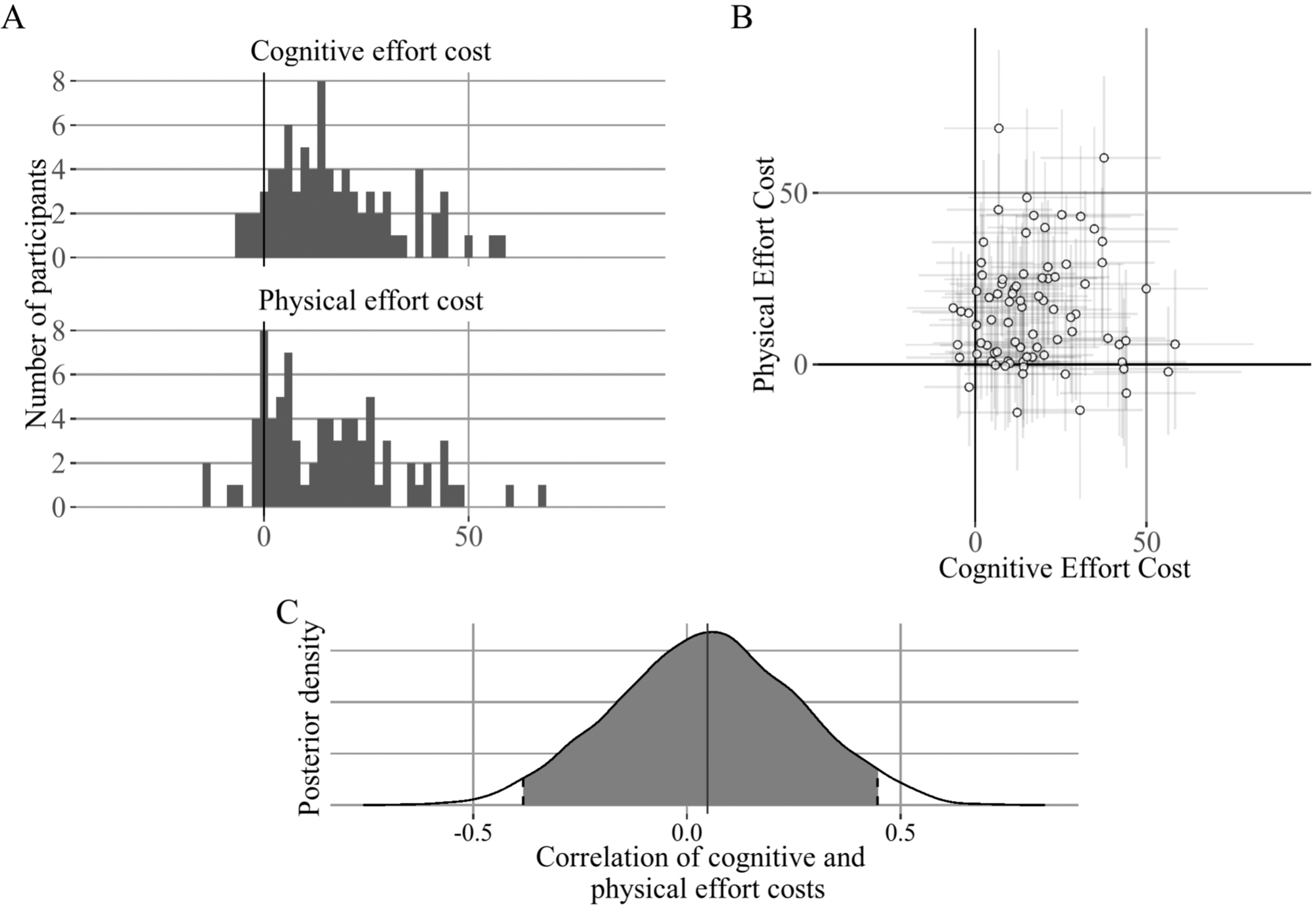
Experiment 2 (N-Back) individual differences in effort costs. A: There were individual differences in cognitive (3-Back) and physical high effort cost. Some participants were insensitive to the manipulation (cost near zero). B: cognitive versus physical effort cost relationship, error bars represent 80% HDI. C: posterior distribution of correlation between high effort cost for cognitive and physical effort. No correlation between cognitive and physical effort cost, wide confidence interval (shaded area represents 95% HDI) suggests sample is underpowered to detect a correlation similar in size to Experiment 1.

#### B. Experiment 3 (Richness)

##### B.1. Methods

In Experiment 3 (Richness) we conducted a study manipulating the tree richness as a benchmark of how participants adjust their exit threshold in response to reward rate (richness was not manipulated in Experiments 1 and 2). We compared two levels of reward richness by adjusting the mean of a normal distribution used to draw the initial reward paid out by a tree. In the ‘lean condition’ the initial reward mean was 15 apples *N* (15, 1) and in the ‘rich condition’ the initial reward mean was 20 apples *N* (20, 1). We tested all combinations of the effort and richness orchard types and counterbalanced block order within effort type. We predicted participants would lower their threshold (exit later) in the lean condition because reward rate is lower in the lean compared to the rich condition. This would confirm that participants still adhere to predictions of the Marginal Value Theorem even in our novel experiment context where effort was added to the travel.

The richness manipulation was conducted during piloting studies of physical effort version of the Effort Foraging Task. There were several differences between the pilot studies (Experiment 3) and the main experiments (1 and 2). Pilot studies were conducted in the laboratory (rather than remotely). For pilot studies we pre-screened participants to have relatively low Need for Cognition (we did not do so in Experiments 1, 2, or 4). The pre-screen survey was completed online no later than 24 hours before the study. Participants gave written consent to complete the pre-screen. To avoid explicit cueing of the objective of the study we administered two foil self-report scales following the Need for Cognition scale: the Individualism and Collectivism Scale (Triandis & Gelfand, 1998) and the Ambiguity Tolerance Scale (Mac Donald, 1970). Participants with Need for Cognition scores less than or equal to 70 points (out of 90 possible points) were invited to the study. Participants again gave written consent to participate in the study.

43 participants volunteered for Experiment 3 (Richness) (24 female, 19 male, 18-34 years old, mean age = 21.5 years *±* 3.7). Experiment 3 (Richness) includes two pilot studies in which richness was manipulated. Because of the heterogeneity of methods, we did not fit the MVT model or estimate the relationship between cognitive and physical effort cost in Experiment 3. In the ‘button-pressing rate’ version, participants had to maintain a fixed rate (smaller number of presses per second vs. larger number of presses per second). In the ‘button-press count’ version, participants had to complete a smaller number or larger number of their maximum calibrated presses (this was the same physical effort requirement as in the main experiments). The cognitive effort requirement in both versions was the N-Back task (same as in Experiment 2 (N-Back)). There were 21 participants in button-pressing rate version and 19 participants in button-pressing count version. Three participants were excluded due to poor button pressing performance (> 2SD uncompleted presses). In both versions participants completed 4 N-Back blocks followed by 4 rapid button pressing blocks. Orchard duration was 5 minutes in the button-press rate version, and 7 minutes in the button-press count version. Harvest and travel time were the same for Experiments 1 (N-Back) and 3 (Richness).

##### B.2. Analysis methods

To test whether participants responded as predicted to the richness manipulation, we fitted a mixed-effects linear regression model to exit thresholds (using the lme4 package in the R language, Bates et al., 2022). The model predicted exit threshold (expected (log) apples) by orchard type separately fit for all conditions (for cognitive high and low effort, and physical high and low effort, and scarce and rich orchards) for all participants. Then we computed a multi degrees-of-freedom test on the linear mixed-effects model (using contestMD function of the lmerTest package (Kuznetsova et al., 2020)). The contrast tested whether the mean-value parameters are significantly different in the scarce compared to the rich condition (collapsing over all the different travel tasks).

#### C. Experiment 4 (Instruct fixed travel time)

##### C.1. Methods

To evaluate whether subjective time may have been a confounding driver of effort avoidance in this task, we conducted an additional ‘Experiment 4’, in which we explicitly instructed participants that the travel time was fixed between effort levels and quizzed them on this fact. This was a replication of Experiment 1 with the only difference being the addition of explicit instructions that the travel time was fixed across all conditions. 71 Prolific participants volunteered for the study (37 females, 30 males, 4 prefer not to answer, mean age 36.27 years *±* 10.56, range 19-62 years). We applied the exclusion cutoffs from Experiment 1 (Tab. S8) resulting in 54 participants included in analyses.

The instructions added were:

‘The time it takes to get to a new tree is fixed. It is NOT RELATED to which orchard you are in or whether you are completing the [matching/small presses] or [mismatching/large presses] trials.’

‘You will visit 4 orchards and spend 16 minutes total. Your choices in the game will NOT make the experiment end earlier. Remember, every apple you harvest earns you money! You can earn up to $2 harvesting the Oddball Numbers Apples Game orchards.’

We also added a quiz question that participants needed to answer correctly (correct answer in bold) to begin the experiment:

‘Question: What determines how long it takes to travel to a new tree?

Options:

The background color.

The time is fixed.

Whether you are completing matching or mismatching trials of the Oddball Number Game./Whether you are completing the small or large presses of the Button Pressing Game.’

##### C.2. Analysis methods

To test whether participants still changed their threshold in the high relative to low effort task, we fitted a mixed-effects linear regression model to exit thresholds (using the lme4 package in the R language, Bates et al., 2022). The model predicted exit threshold (expected (log) apples) by orchard type separately fit for all conditions (for cognitive high and low effort, and physical high and low effort) for all participants. Then we computed a multi degrees-of-freedom test on the linear mixed-effects model (using the contestMD function of the lmerTest package, Kuznetsova et al., 2020). We found a significant decrease in exit threshold, replicating Experiment 1 even when participants were explicitly instructed the travel time is fixed (linear mixed-effects regression estimate for MSIT (Interference-Congruent):= −0.318 apples, df=49.50, F=12.66, p<0.001, physical (smaller-larger) = −0.391, df=47.72, F=5.66, p<0.021, Fig. S9).

**Fig. S9.**
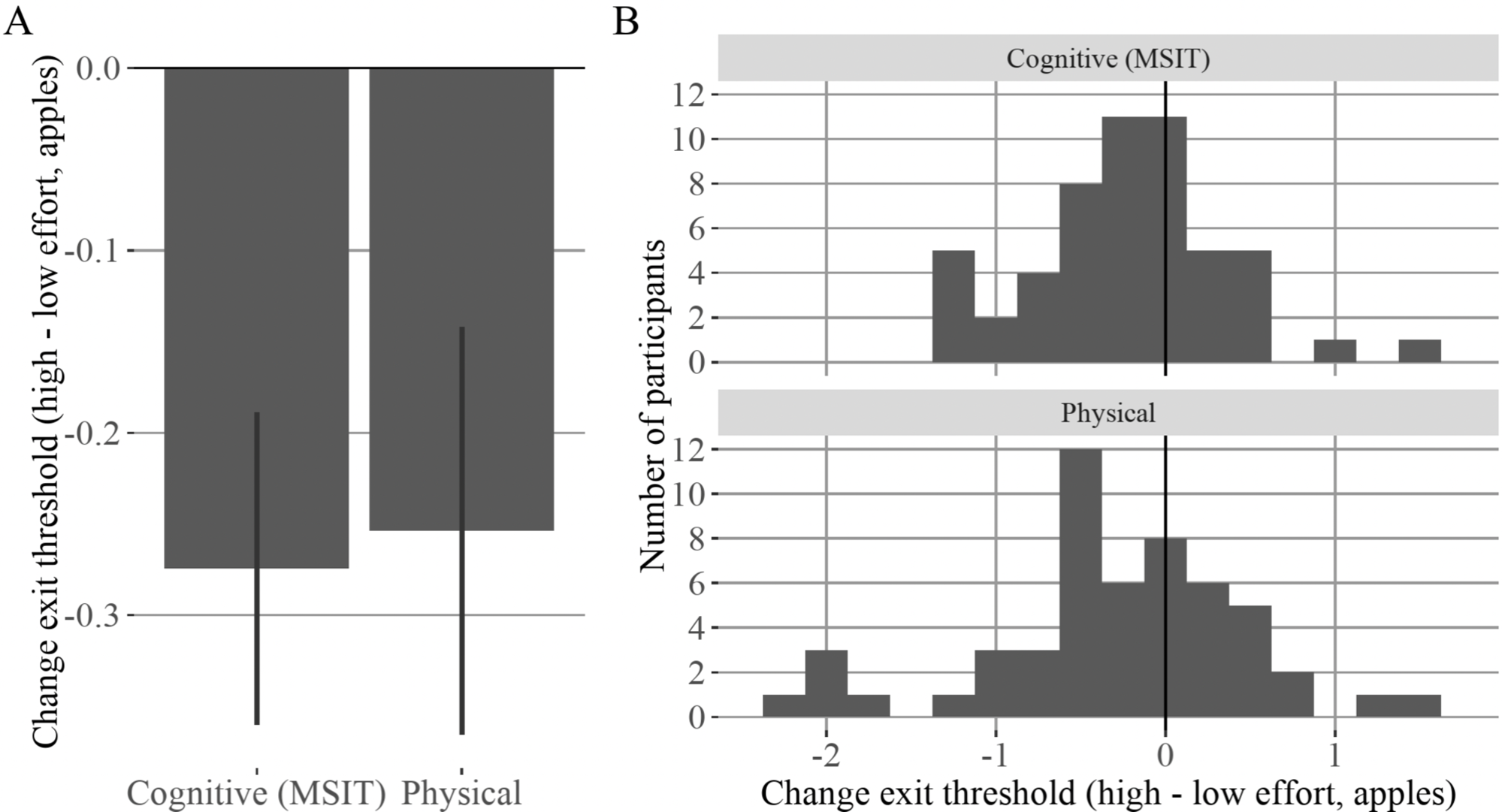
Experiment 4 (Instruct fixed time), participants avoid effort when explicitly instructed that the travel time is fixed across all conditions. A: y-axis indicates change in exit threshold for cognitive and physical effort (high - low effort level), x-axis indicates effort type, B: histogram of individual differences in change in exit threshold by effort condition, top row cognitive effort, bottom row physical effort.

**Table S5.** Foraging environment parameters comparison chart. Column 1: environment parameter for; column 2: Experiments 1 & 4 (MSIT), column 3: Experiment 2 (N-Back), column 4: Experiment 3 (Richness) Experiment 1 (MSIT). Third column: Experiment 2 (N-Back).

| Parameter | Experiments 1 & 4 (MSIT) | Experiment 2 (N-Back) | Experiment 3 (Richness) |
| --- | --- | --- | --- |
| Harvest time | 2 seconds | 2 seconds | 2 seconds |
| Travel time | 8.33 seconds | 20 seconds | 20 seconds |
| Cognitive Travel Tasks | MSIT Congruent, Interference | 1-Back, 3-Back | 1-Back, 3-Back |
| Physical Travel Tasks | Larger (100% max), Smaller (50%) number of presses | (100% and 50% max) | Number or rate of presses |
| Block duration | 4 minutes | 7 minutes | 7 minutes |
| Number blocks per condition | 2 | 2 | 2 |
| Initial reward | $\mathcal{N}(15, 1)$ | $\mathcal{N}(15, 1)$ | $\mathcal{N}(15, 1), \mathcal{N}(20, 1)$ |
| Decay rate | $\beta(14.13, 2.03)$ | $\beta(14.13, 2.03)$ | $\beta(14.13, 2.03)$ |
| Total Number of blocks | 8 | 8 | 8 |

**Table S6.** Experiment 1 Race and Hispanic or Latino ethnicity. Participants reported their race and Hispanic or Latino ethnicity from a multiple choice table. The left column indicates their first selection, the middle column indicates their second selection (if any). The right column indicates the number of participants.

| **Race** | **Number of participants** |
| --- | --- |
| American Indian/Alaskan Native | 2 |
| Asian | 22 |
| Asian & White | 2 |
| Black or African American | 55 |
| Black or African American & Asian | 1 |
| Black or African American & White | 3 |
| White | 516 |
| White & More than one race | 9 |
| More than one race | 36 |
| Prefer not to answer | 32 |
| **Hispanic or Latino** |  |
| Hispanic or Latino | 119 |
| Not Hispanic or Latino | 543 |
| Prefer not to answer | 16 |

### 6. Simulation to find best threshold

We simulated the best foraging threshold by creating a foraging environment with an agent with a fixed exit threshold and observing the resulting reward rate. We used a policy iteration algorithm to find the maximal reward rate for a given foraging environment. The foraging environment was defined by the following parameters from our experiments; the harvest time (2 seconds), travel time (8.33 seconds), the distribution of initial rewards to a tree *N* (15, 1) distribution of the decay function (beta distribution, *β*(14.90873, 2.033008)). We assumed the agent knew the mean depletion rate (0.88 multiplied by the previous reward) and used this value to predict the expected reward on the current trial. If the predicted reward was less than or equal to the agent’s threshold it exited the patch *R_e_ ≤ ρ*, otherwise it harvested the patch which yielded reward. We simulated 840 ‘seconds’ of foraging time for all experiments (though the result should be robust to duration). The simulation outputs were the ‘best threshold’ (threshold that yielded the highest reward rate, results vary slightly by simulation run), the resulting ‘best reward rate’, as well as the mean and standard deviation number of harvests to reach that exit threshold.

The agents’ threshold parameter was initialized at 4 apples. For an iteration i, the threshold was set as the mean reward rate observed in iteration i-1, this allowed the threshold to gradually improve in terms of reward rate between iterations. The simulation stopped and the best threshold was determined based on the stopping threshold of a 0.001 apple per second improvement in reward rate on iteration i compared iteration i-1 (with a maximum of 200 iterations). Best exit threshold policy in simulated data (not including effort costs) was 6.78 apples, the reward rate achieved with best threshold was 3.39 apples per second, and on average it took 6.77 *±* 1.69 harvests to reach the best threshold (Tab. S7).

**Table S7.** Best exit threshold policy in simulated data. Column 1: best exit threshold policy parameters; column 2: Experiments 1 & 4 (MSIT); column 3: Experiment 2 (N-Back) and Experiment 3 (Scarce condition); column 4 (Rich condition). Row 1: best threshold from simulation (apples); row 2: reward rate achieved with best threshold (apples per second); row 3: mean harvests it took to reach the best threshold; row 4: standard deviation of harvests it took to reach the best threshold.

|  | Experiment 1 & 4 (MSIT) | Experiment 2 (N-Back) & Experiment 3 (Scarce) | Experiment 3 (Rich) |
| --- | --- | --- | --- |
| Best threshold (apples) | 6.78 | 4.65 | 6.17 |
| Reward rate (apples per second) | 3.39 | 2.32 | 2.37 |
| Number of harvests $\mu$ | 6.77 | 9.95 | 9 |
| Number of harvests $SD$ | 1.69 | 2.01 | 2.56 |

### 7. Exclusions Experiment 1

Experiment 1 participants were excluded if they missed many harvest trials (if they did not respond after 1 second deadline in greater than 10% of all harvest trials). Participants were excluded if they performed poorly on any of the travel tasks (using the metrics MSIT congruent and interference trial error rate, percent smaller presses uncompleted, percent larger presses uncompleted). For each travel task we computed the group mean and standard deviation, and excluded participants who were 2 standard deviations below the group mean performance (Tab. S8).

**Table S8.** Effort Foraging Task Behavior Based Exclusions Experiment 1. Column 1: basis of exclusion, column 2: number of participants, column 3: exclusion cutoff value, column 4: group mean and standard deviation. Number of participants outliers by exclusion criteria. Participants could be excluded on multiple grounds; therefore, the number of outlier participants does not add up to the number of total number of excluded participants.

| Exclusion criteria | N outliers | Cutoff | Mean, SD |
| --- | --- | --- | --- |
| Did not complete experiment | 5 |  |  |
| Foraging trial response deadline missed (% trials) | 30 | 10 | 2.8, 5.9 |
| MSIT Congruent error rate (> 2SD) | 10 | 16.6 | 2.7, 7.0 |
| MSIT Interference error rate (> 2SD) | 51 | 54.2 | 19.2, 17.5 |
| Smaller number of presses uncompleted (%,>2SD) | 12 | 14.1 | 1.9, 6.1 |
| Larger number of presses uncompleted (%,>2SD) | 29 | 21.0 | 4.9, 8.0 |
| Number of exit trials per condition (<2SD) | 11 | 5.4 | 18.6, 6.6 |
| Change exit threshold cognitive (<2SD \| >2SD) | 30 | <-2.52 \| >1.91 | -0.31, 1.11 |
| Change exit threshold physical (<2SD \| >2SD) | 31 | <-2.77 \| >2.0 | -0.39, 1.19 |
| Initial Sample Size | 678 |  |  |
| Final Sample Size | 537 |  |  |

### 8. Foraging task training

In Experiment 1 the task began with training the travel task for the first effort cost variant for a particular participant (this could be the cognitive or physical effort task). Next came instructions for the foraging task in general (without mentioning the effortful travel requirement), and participants completed a practice block (90 seconds) of the foraging task with no travel task. Then participants were instructed that they would have to complete the effortful travel task when traveling, and they completed two practice blocks (one per effort level, 90 seconds each). Then, participants completed the main foraging task for the first travel task type (4 blocks, 4 minutes per block, with self-paced breaks between blocks). After completing all the blocks of the first travel task, participants began training on the second travel task. They were instructed that they would continue to play the foraging task, but the travel task had changed. They practiced the foraging task with the second travel task type (one practice block per effort level, 90 seconds each). Finally, they completed the main foraging task for the second travel task type (4 blocks, 4 minutes per block).

### 9. Rapid key-pressing task training

In Experiment 1 key-press training began with a calibration phase (three rounds) to determine the maximum number of presses participants were able to complete in the travel time (7.5 seconds of effort task time). A counter was displayed on the center of the screen showing how many presses a participant had made. The instructions suggested participants were being compared to others, and encouraged them to press as fast as possible, each round they were encouraged to press faster than they had the previous rounds (see instructions in SI Text 10). Then we used each participant’s mean number of presses across rounds as their ‘maximum number’. We enforced a minimum ‘maximum number’ value of 15 presses. The Larger Number of Presses condition tasked participants with completing 100% of their maximum, and the Smaller Number of Presses condition tasked participants with completing 50% of their maximum. Participants were told that there was a larger and smaller number, but not what that number was or how it was determined. Then participants practiced a single effort level. Effort level order was counterbalanced. Practice for an effort level began with a single mini-block the duration of the foraging travel time. Then participants had to complete 5 mini-blocks reaching the required number of presses to move on. This was meant to establish the expectation that participants would perform well on the travel task, even though there were no incentives or punishments associated with travel task performance during the foraging task.

### 10. Foraging task instructions

“Welcome to the experiment! Thank you for participating. This experiment will require you to press buttons on your keyboard repeatedly, applying varying amounts of physical effort. If you have any history of any sort of hand injury or pain with typing (e.g., which could make either fast button pressing or stretching your hand uncomfortable) please do not complete this task. You must wait a minimum of 5 seconds before you are able to progress to a new slide of instructions. You will know you can click to a new slide when the “Next” button changes.”

“Welcome to the Apples Game! For your completion of this task, you will receive a potential bonus between $0 and $5. Please read the instructions carefully. There will be a quiz at the end of these instructions to check your understanding. In this game, you will make choices that earn you money. Imagine you are a farmer, and you are harvesting apples from trees in your multiple orchards. On every trial within an orchard, you will see a tree: To HARVEST the tree, press the down arrow key with your right hand. Do this when the circle below the tree is white. When you harvest the tree it gives you apples. These apples are worth real money that you will earn on top of the money for participating in this study. Now, try harvesting the tree three times in a row. Press the HARVEST key to collect apples from the tree. [Press “Next” to practice using the HARVEST key 3x]”

“The more times you harvest a tree, the fewer apples it gives you! On any trial, instead of accepting the number of apples the tree is giving you, you have the option to TRAVEL to a new tree. To TRAVEL to a new tree press the right arrow key with your right hand. Do this when the circle below the tree is white. Now, harvest the tree once and travel from one tree to another. Do this three times. Press the TRAVEL key to move to a new tree. [Press “Next” to practice using the TRAVEL key 3x]”

“Different trees give you different number of apples at the start. Exactly how many apples a tree starts with changes from tree to tree. The starting number of apples for a tree is NOT RELATED to how the tree looks or which orchard you are in. HARVESTING takes some time but earns you apples. TRAVELING takes longer, and you cannot harvest apples during traveling. But it brings you to a new tree with a full supply. You have to decide how to spend your limited time in an orchard – harvesting or traveling. You have to HARVEST each new tree once before traveling away from it. If you take too long to make a choice you will miss a turn and see this message. The more turns you miss, the less time you have to harvest apples, and you will earn less apples. That is most of what you have to know to be a great farmer. Let’s go through a short practice orchard in the Apples Game.”

Then participants completed a 1.5-minute orchard with no travel task. Then the learned one of the effortful travel tasks (order of effort types counterbalanced).

#### A. Rapid keypressing task training instructions

To train the button-pressing task participants were taught the button-press hold keys and then we challenged them to press as fast as they could across three calibration blocks. “You will now play the Button Pressing Game. Please read the instructions carefully. To play the Button Pressing Game, hold down the hold keys while rapidly pressing the press key. For your left hand, put your pinky on the A key, ring finger on the S key, middle finger on the D key, and pointer finger on the V key. For your right hand, put your pointer finger on the N key and your ring finger on the L key. First, let’s try getting as many button presses as we can.”

You will practice by playing 3 sets of button presses.

Great job completing those button presses! Now you will complete more presses.

Calibration block 1: “We want to know how fast you can press a button compared to other people. The computer will count up the number of presses you can complete for a short block. With your left pinky finger press the ‘A’ key as many times as you can before the block ends. You can stop pressing when the display changes to ‘Complete!’ “When you are ready to begin, start pressing the ‘A’ key fast as you can!”

Calibration block 2: “Great job! Take a break. When you are ready you will complete another short block. Try and complete even more presses this block! With your left pinky finger press the ‘A’ key as many times as you can before the block ends. You can stop pressing when the display changes to ‘Complete!’ When you are ready to begin, start pressing the ‘A’ key fast as you can!”

Calibration block 3: “Great job! Take a break. When you are ready you will complete ONE FINAL short block. Think you can complete any more presses? With your left pinky finger press the ‘A’ key as many times as you can before the block ends. You can stop pressing when the display changes to ‘Complete!’ When you are ready to begin, start pressing the ‘A’ key fast as you can!”

After this we computed the maximum number of presses they completed in the calibration round and set their larger (100% max) and smaller (50% max) button presses requirement.

“In this experiment sometimes you will complete a LARGE number of presses, and other times you will complete a SMALL number. You will now practice completing the [SMALL/LARGE] number of button presses. Press the ‘A’ key until you reach the goal number of times shown on the screen. If you do not finish in time, you will see a black circle at the end of the block. Don’t worry about finishing in time; just focus on learning the timing. When you are ready to begin, start pressing the ‘A’ key.”

“You will now practice completing the [SMALL/LARGE] number of button presses. Press the ‘A’ key until you reach the goal number of times shown on the screen. If you do not finish in time, you will see a black circle at the end of the block. Don’t worry about finishing in time; just focus on learning the timing. When you are ready to begin, start pressing the ‘A’ key.”

Then participants completed mini-blocks. The purpose of these blocks was to establish the expectation of completing the task during the travel time. “You will complete the [SMALL/LARGE] number of presses for [N] miniblocks. Press the F key the number of times shown on the screen. To move on to the next miniblock, you must reach the goal number of presses displayed on the screen before the deadline. If you do not finish in time, you will see a black circle at the end of the block. When you are ready to begin, press the spacebar THEN press the hold keys.”

If participants failed they had to repeat the mini-block:

“Block [X] of [N]. You must repeat this block. Complete the [SMALL/LARGE] number of presses. When you are ready to begin, press the spacebar THEN press the hold keys.” If they succeeded they moved onto the next mini-block:

“Block [X] of [N]. Moving to next block. Complete the [SMALL/LARGE] number of presses. When you are ready to begin, press the spacebar THEN press the hold keys.”

#### B. Physical Effort Foraging Task Instructions

“Here are some things to keep in mind: Sometimes you will complete a small number of presses and the background will be blue. Other times, you will complete a large number of presses and the background will be orange. If you complete the presses before the period is up, you will see “Complete!” on your screen. If you press fewer than the number of required presses, you will see a black circle. Please try to avoid seeing this circle. Now let’s practice some more button presses! From now, you will play the Apples Button Pressing Game to TRAVEL from tree to tree. In some orchards you will complete the small presses when you travel and during those orchards the background will always be blue. In other orchards you will complete the large presses trials when you travel and during those orchards the background will always be orange. If you complete the presses before the travel period is up, you will see “Complete!” on your screen. Please try your best on each trial. If you make 2 errors in a row, including missing trials, you will see a black dot:

There will be a message at the start of each orchard telling you which traveling game you will play. During an orchard if you forget which task you should perform, you can look to the background color when traveling to remember. That is most of what you have to know to be a great farmer. Let’s go through a short practice of the Apples Button Pressing Game.”

Then participants completed a 1.5 minute orchard in the low effort (smaller number of presses) and a 1.5 minute orchard in the high effort (larger number of presses) condition.

Participants were also instructed to take a break after completing half the orchards: “You are finished with the first half, take a break and proceed to the next orchard when you are ready. [Press Spacebar to continue]”

#### C. Cognitive Effort Foraging instructions

“Great job harvesting in your orchard! From now, you will play the Apples Letter Matching Game to TRAVEL from tree to tree. In some orchards you will complete the 1-Back trials when you travel and during those orchards the background will always be blue. In other orchards you will complete the 3-Back trials when you travel and during those orchards the background will always be orange. Please try your best on each trial. If you make 2 errors in a row, including missing trials, you will see a black dot: There will be a message at the start of each orchard telling you which traveling game you will play. During an orchard if you forget which task you should perform you can look to the background color when traveling to remember. That is most of what you have to know to be a great farmer. Let’s go through a short practice of the Apples Letter Matching Game.”

Then participants completed a 1.5-minute orchard in the low effort (1-Back / congruent trials) and a 1.5-minute orchard in the high effort (3-Back / interference trials) condition.

“Great job harvesting apples! Here are a few more things you should keep in mind: You will visit 4 orchards and spend 7 minutes in each. You will know you are being moved to a new orchard when the background color changes, and you see a message like this: ‘You are entering a 1-Back orchard! To earn as many apples as possible, please pay attention to how many apples a tree produces, how its supply of apples decreases, and how long it takes to travel to a new tree! Please press the spacebar to continue to the orchard.’ This is everything you need to know to be a great farmer. You can make up to $5 across these 4 orchards. Remember, every apple you harvest earns you money!”

“Before you begin, you will need to pass a quiz on the instructions. Answering a question wrong will require you to re-read the instructions. Press NEXT to begin the quiz.”

#### D. Foraging comprehension quiz

Before starting the first foraging orchard, participants were given a short quiz on the harvest and travel keys, travel task, and timing per orchard (correct answer in bold).

*Quiz preamble*: “Please answer every question. Answering incorrectly will require you return to the beginning of the instructions.”

*Quiz question 1* : “What determines how long you will spend in each orchard?”.

Options:

‘The number of times you harvest.’

‘The number of times you travel.’

‘The time is fixed (1 minute).’

**‘The time is fixed (7 minutes).’**

*Quiz question 2* : “When you are at a tree, how do you collect apples?”.

Options:

‘Repeatedly pressing the TRAVEL key (right arrow key).’

‘Repeatedly pressing the HARVEST key (the down arrow key).’

‘Pressing the TRAVEL key (right arrow key), once per harvest.’

**‘Pressing the HARVEST key (the down arrow key), once per harvest.’**

*Quiz question 3* : “What happens to the number of apples a tree provides over time?”.

Options:

**‘The number of apples the tree gives you decreases with each harvest.’**

‘The number of apples the tree gives you increases with each harvest.’

’The number of apples the tree gives you does not change with each harvest.’

*Quiz question 4* : “When you are at a tree, how do you leave the tree?”.

Options:

‘Repeatedly pressing the TRAVEL key (right arrow key).’

‘Repeatedly pressing the HARVEST key (the down arrow key).’

**‘Pressing the TRAVEL key (right arrow key), once.’**

‘Pressing the HARVEST key (the down arrow key), once.’

*Quiz question 5* (this question depends on the travel task): “What factor makes orchards differ from one another?”.

If the travel task was N-Back:

‘How any apples a tree gives on the first harvest.’

‘The rate at which the number of apples at a tree falls.’

**‘In some orchards you will play 1-Back trials of the Letter Matching Game. In other orchards you will play 3-Back trials of the game.’**

If the travel task was the Multi-source Interference Task:

‘How any apples a tree gives on the first harvest.’

’The rate at which the number of apples at a tree falls.’

**‘In some orchards you will play matching trials of the Oddball Number Game. In other orchards you will play mismatching trials of the game.’**

If the travel task was rapid keypressing:

’How any apples a tree gives on the first harvest.’

‘The rate at which the number of apples at a tree falls.’

**‘In some orchards you will play small presses trials of the Button Pressing Game. In other orchards you will play large presses trials of the game.’**

*Quiz question 6* : “What determines your bonus?”.

Options:

**‘Total number of apples collected over the entire experiment.’**

‘Total number of apples collected on a randomly selected orchard.’

‘Highest single apples value received on a randomly selected orchard.’

#### E. Task debrief survey

Free response questions from Experiment 2 (N-Back) including subjective awareness of travel effort cost manipulation:

How did you decide when to leave a tree?

Did the number of letters back in the traveling task change your decision to travel to a new tree? If yes, how so?

Did the number of button presses in the traveling task change your decision to travel to a new tree? If yes, how so?

Did the duration of the travel change your decision to travel to a new tree? If yes, how so?

What strategies did you use when you were harvesting a tree?

What strategies did you use in the letter matching task?

What strategies did you use in the button pressing task?

Travel task effort ratings (question order shuffled) [1 - Not effortful at all, 2, 3, 4, 5 - Very effortful]:

How effortful did you find responding 3 letters back?

How effortful did you find responding 1 letter back?

How effortful did you find completing the LARGE number of presses?

How effortful did you find completing the SMALL number of presses?

Travel task enjoyment ratings (question order shuffled) [1 - Not enjoyable at all, 2, 3, 4, 5 - Very enjoyable]:

How enjoyable did you find responding 1 letter back?

How enjoyable did you find completing the SMALL number of presses?

How enjoyable did you find responding 3 letters back?

How enjoyable did you find completing the LARGE number of presses?

Boredom ratings (question order shuffled) [1 - Not bored at all, 2, 3, 4, 5 - Extremely bored]:

How bored were you during orchards for which the traveling task was to respond 3 letters back?

How bored were you during orchards for which the traveling task was to respond 1 letter back?

How bored were you during orchards for which the traveling task was to complete the LARGE number of presses?

How bored were you during orchards for which the traveling task was to complete the SMALL number of presses?

Did you always use your non-dominant pinky finger when completing the button pressing task? [yes or no]

Subjective time perception of travel [slider 0 to 60 seconds, increments of 1 second]:

How long do you think the travel time was during orchards for which the traveling task was to respond 3 letters back?

How long do you think the travel time was during orchards for which the traveling task was to respond 1 letter back?

How long do you think the travel time was during orchards for which the traveling task was to complete the LARGE number of presses?

How long do you think the travel time was during orchards for which the traveling task was to complete the SMALL number of presses?

## Notes

https://osf.io/a4r2e/

### Competing Interest Statement

The authors have declared no competing interest.

### Summary of Updates

Manuscript revised based on peer reviewer feedback, and is in resubmission. New data and analyses were added, and text was updated. A link to open access materials were added. The canonical correlation analysis (CCA) results relating task parameters to self-reported symptoms were changed, after correcting an error we discovered during the revision. We inadvertently computed degrees of freedom for the CCA based on the total number of participants (N=678, a number which is manually input to the statistical test). In addition, the data we used for the CCA contained the full group who had task behavior (N=537, including those for whom we did not have self-report data, who were inappropriate for inclusion in the CCA). We corrected this error re-ran the full CCA analyses and associated statistics using only the 430 participants for whom we have full datasets, and these results are presented in the revised manuscript.

## References

Wilks, S. S. (1935). On the independence of k sets of normally distributed statistical variables [Publisher: [Wiley, Econometric Society]]. Econometrica, 3 (3), 309–326. 10.2307/1905324

Neisser, U. (1967). Cognitive psychology. Appleton-Century-Crofts.

Charnov, E. L. (1976). Optimal foraging, the marginal value theorem. Theoretical Population Biology, 9 (2), 129–136. 10.1016/0040-5809(76)90040-X

Cacioppo, J. T., Petty, R. E., & Kao, C. F. (1984). The efficient assessment of need for cognition. Journal of Personality Assessment, 48 (3), 306–307. 10.1207/s15327752jpa480313

Stephens, D. W., & Krebs, J. R. (1986). Foraging theory (Vol. 1). Princeton University Press. 10.2307/j.ctvs32s6b

Carver, C. S., & White, T. L. (1994). Behavioral inhibition, behavioral activation, and affective responses to impending reward and punishment: The BIS/BAS scales [Place: US Publisher: American Psychological Association]. Journal of Personality and Social Psychology, 67 (2), 319–333. 10.1037/0022-3514.67.2.319

Snaith, R. P., Hamilton, M., Morley, S., Humayan, A., Hargreaves, D., & Trigwell, P. (1995). A scale for the assessment of hedonic tone the snaith-hamilton pleasure scale. The British Journal of Psychiatry: The Journal of Mental Science, 167 (1), 99–103. 10.1192/bjp.167.1.99

Baumeister, R. F., & Heatherton, T. F. (1996). Self-regulation failure: An overview [Place: US Publisher: Lawrence Erlbaum]. Psychological Inquiry, 7 (1), 1–15. 10.1207/s15327965pli07011

Pirolli, P., & Card, S. (1999). Information foraging [Place: US Publisher: American Psychological Association]. Psychological Review, 106, 643–675. 10.1037/0033-295X.106.4.643

Nystrom, L. E., Braver, T. S., Sabb, F. W., Delgado, M. R., Noll, D. C., & Cohen, J. D. (2000). Working memory for letters, shapes, and locations: fMRI evidence against stimulus-based regional organization in human prefrontal cortex. NeuroImage, 11 (5), 424–446. 10.1006/nimg.2000.0572

Kroenke, K., Spitzer, R. L., & Williams, J. B. (2001). The PHQ-9: Validity of a brief depression severity measure. Journal of General Internal Medicine, 16 (9), 606–613. 10.1046/j.1525-1497.2001.016009606.x

Bush, G., & Shin, L. M. (2006). The multi-source interference task: An fMRI task that reliably activates the cingulo-frontal-parietal cognitive/attention network. Nature Protocols, 1 (1), 308–313. 10.1038/nprot.2006.48

Khan, A., Sharma, N. K., & Dixit, S. (2006). Effect of cognitive load and paradigm on time perception [Place: India Publisher: Indian Academy of Applied Psychology]. Journal of the Indian Academy of Applied Psychology, 32 (1), 37–42.

Spitzer, R. L., Kroenke, K., Williams, J. B. W., & Löwe, B. (2006). A brief measure for assessing generalized anxiety disorder: The GAD-7. Archives of Internal Medicine, 166 (10), 1092–1097. 10.1001/archinte.166.10.1092

Cella, D., Yount, S., Rothrock, N., Gershon, R., Cook, K., Reeve, B., Ader, D., Fries, J. F., Bruce, B., & Rose, M. (2007). The patient-reported outcomes measurement information system (PROMIS). Medical care, 45 (5), S3–S11. 10.1097/01.mlr.0000258615.42478.55

Evans, D. E., & Rothbart, M. K. (2007). Developing a model for adult temperament. Journal of Research in Personality, 41 (4), 868–888. 10.1016/j.jrp.2006.11.002

Niv, Y., Daw, N. D., Joel, D., & Dayan, P. (2007). Tonic dopamine: Opportunity costs and the control of response vigor. Psychopharmacology, 191 (3), 507–520. 10.1007/s00213-006-0502-4

Walton, M. E., Rudebeck, P. H., Bannerman, D. M., & Rushworth, M. F. S. (2007). Calculating the cost of acting in frontal cortex. Annals of the New York Academy of Sciences, 1104, 340–356. 10.1196/annals.1390.009

Hills, T. T., Todd, P. M., & Goldstone, R. L. (2008). Search in external and internal spaces: Evidence for generalized cognitive search processes [Place: United Kingdom Publisher: Wiley-Blackwell Publishing Ltd.]. Psychological Science, 19 (8), 802–808. 10.1111/j.1467-9280.2008.02160.x

Chib, V. S., Rangel, A., Shimojo, S., & O’Doherty, J. P. (2009). Evidence for a common representation of decision values for dissimilar goods in human ventromedial prefrontal cortex. The Journal of Neuroscience, 29 (39), 12315–12320. 10.1523/JNEUROSCI.2575-09.2009

Lewandowski, D., Kurowicka, D., & Joe, H. (2009). Generating random correlation matrices based on vines and extended onion method. Journal of Multivariate Analysis, 100 (9), 1989–2001. 10.1016/j.jmva.2009.04.008

Marcora, S. M., Staiano, W., & Manning, V. (2009). Mental fatigue impairs physical performance in humans. Journal of Applied Physiology (Bethesda, Md.: 1985), 106 (3), 857–864. 10.1152/japplphysiol.91324.2008

Treadway, M. T., Buckholtz, J. W., Schwartzman, A. N., Lambert, W. E., & Zald, D. H. (2009). Worth the ‘EEfRT’? the effort expenditure for rewards task as an objective measure of motivation and anhedonia [Publisher: Public Library of Science]. PLOS ONE, 4 (8), e6598. 10.1371/journal.pone.0006598

Wilke, A., Hutchinson, J. M. C., Todd, P. M., & Czienskowski, U. (2009). Fishing for the right words: Decision rules for human foraging behavior in internal search tasks. Cognitive Science, 33 (3), 497–529. 10.1111/j.1551-6709.2009.01020.x

Block, F., & Gellersen, H. (2010). The impact of cognitive load on the perception of time. Proceedings of the 6th Nordic Conference on Human-Computer Interaction: Extending Boundaries, 607–610. 10.1145/1868914.1868985

Block, R. A., Hancock, P. A., & Zakay, D. (2010). How cognitive load affects duration judgments: A meta-analytic review. Acta Psychologica, 134 (3), 330–343. 10.1016/j.actpsy.2010.03.006

Kool, W., McGuire, J. T., Rosen, Z. B., & Botvinick, M. M. (2010). Decision making and the avoidance of cognitive demand. Journal of Experimental Psychology. General, 139 (4), 665–682. 10.1037/a0020198

Krebs, R. M., Boehler, C. N., & Woldorff, M. G. (2010). The influence of reward associations on conflict processing in the stroop task. Cognition, 117 (3), 341–347. 10.1016/j.cognition.2010.08.018

Cĺery-Melin, M.-L., Schmidt, L., Lafargue, G., Baup, N., Fossati, P., & Pessiglione, M. (2011). Why don’t you try harder? an investigation of effort production in major depression. PloS One, 6 (8), e23178. 10.1371/journal.pone.0023178

Hayden, B. Y., Pearson, J. M., & Platt, M. L. (2011). Neuronal basis of sequential foraging decisions in a patchy environment. Nature Neuroscience, 14 (7), 933–939. 10.1038/nn.2856

Levy, D. J., & Glimcher, P. W. (2011). Comparing apples and oranges: Using reward-specific and reward-general subjective value representation in the brain. The Journal of Neuroscience, 31 (41), 14693–14707. 10.1523/JNEUROSCI.2218-11.2011

Hills, T. T., Jones, M. N., & Todd, P. M. (2012). Optimal foraging in semantic memory. Psychological Review, 119 (2), 431–440. 10.1037/a0027373

Kolling, N., Behrens, T. E. J., Mars, R. B., & Rushworth, M. F. S. (2012). Neural mechanisms of foraging [Publisher: American Association for the Advancement of Science]. Science, 336 (6077), 95–98. 10.1126/science.1216930

Levy, D. J., & Glimcher, P. W. (2012). The root of all value: A neural common currency for choice. Current opinion in neurobiology, 22 (6), 1027–1038. 10.1016/j.conb.2012.06.001

Rigoux, L., & Guigon, E. (2012). A model of reward- and effort-based optimal decision making and motor control. PLoS computational biology, 8 (10), e1002716. 10.1371/journal.pcbi.1002716

Schmidt, L., Lebreton, M., Cĺery-Melin, M.-L., Daunizeau, J., & Pessiglione, M. (2012). Neural mechanisms underlying motivation of mental versus physical effort [Publisher: Public Library of Science]. PLOS Biology, 10 (2), e1001266. 10.1371/journal.pbio.1001266

Todd, P. M., Hills, T. T., & Robbins, T. W. (Eds.). (2012). Cognitive search: Evolution, algorithms, and the brain. The MIT Press. 10.7551/mitpress/9780262018098.001.0001

Treadway, M. T., Bossaller, N., Shelton, R. C., & Zald, D. H. (2012). Effort-based decision-making in major depressive disorder: A translational model of motivational anhedonia. Journal of abnormal psychology, 121 (3), 553–558. 10.1037/a0028813

Treadway, M. T., Buckholtz, J. W., Cowan, R. L., Woodward, N. D., Li, R., Ansari, M. S., Baldwin, R. M., Schwartzman, A. N., Kessler, R. M., & Zald, D. H. (2012). Dopaminergic mechanisms of individual differences in human effort-based decision-making. The Journal of Neuroscience, 32 (18), 6170–6176. 10.1523/JNEUROSCI.6459-11.2012

Kurzban, R., Duckworth, A., Kable, J. W., & Myers, J. (2013). An opportunity cost model of subjective effort and task performance. The Behavioral and brain sciences, 36 (6), 10.1017/S0140525X12003196. 10.1017/S0140525X12003196

Shenhav, A., Botvinick, M. M., & Cohen, J. D. (2013). The expected value of control: An integrative theory of anterior cingulate cortex function. Neuron, 79 (2), 217–240. 10.1016/j.neuron.2013.07.007

Westbrook, A., Kester, D., & Braver, T. S. (2013). What is the subjective cost of cognitive effort? load, trait, and aging effects revealed by economic preference. PloS One, 8 (7), e68210. 10.1371/journal.pone.0068210

Wolfe, J. M. (2013). When is it time to move to the next raspberry bush? foraging rules in human visual search. Journal of Vision, 13 (3), 10. 10.1167/13.3.10

Barch, D. M., Treadway, M. T., & Schoen, N. (2014). Effort, anhedonia, and function in schizophrenia: Reduced effort allocation predicts amotivation and functional impairment. Journal of Abnormal Psychology, 123 (2), 387–397. 10.1037/a0036299

Blanchard, T. C., & Hayden, B. Y. (2014). Neurons in dorsal anterior cingulate cortex signal postdecisional variables in a foraging task [Publisher: Society for Neuroscience Section: Articles]. Journal of Neuroscience, 34 (2), 646–655. 10.1523/JNEUROSCI.3151-13.2014

Wilson, R. C., Geana, A., White, J. M., Ludvig, E. A., & Cohen, J. D. (2014). Humans use directed and random exploration to solve the explore-exploit dilemma. Journal of Experimental Psychology. General, 143 (6), 2074–2081. 10.1037/a0038199

Wolf, D. H., Satterthwaite, T. D., Kantrowitz, J. J., Katchmar, N., Vandekar, L., Elliott, M. A., & Ruparel, K. (2014). Amotivation in schizophrenia: Integrated assessment with behavioral, clinical, and imaging measures. Schizophrenia Bulletin, 40 (6), 1328–1337. 10.1093/schbul/sbu026

Yang, X.-H., Huang, J., Zhu, C.-Y., Wang, Y.-F., Cheung, E. F. C., Chan, R. C. K., & Xie, G.-R. (2014). Motivational deficits in effort-based decision making in individuals with subsyndromal depression, first-episode and remitted depression patients. Psychiatry Research, 220 (3), 874–882. 10.1016/j.psychres.2014.08.056

Blanchard, T. C., & Hayden, B. Y. (2015). Monkeys are more patient in a foraging task than in a standard intertemporal choice task [Publisher: Public Library of Science]. PLOS ONE, 10 (2), e0117057. 10.1371/journal.pone.0117057

Carter, E. C., Pedersen, E. J., & McCullough, M. E. (2015). Reassessing intertemporal choice: Human decision-making is more optimal in a foraging task than in a self-control task. Frontiers in Psychology, 6, 95. 10.3389/fpsyg.2015.00095

de Leeuw, J. R. (2015). jsPsych: A JavaScript library for creating behavioral experiments in a web browser. Behavior Research Methods, 47 (1), 1–12. 10.3758/s13428-014-0458-y

Hills, T. T., Todd, P. M., Lazer, D., Redish, A. D., Couzin, I. D., & Group, t. C. S. R. (2015). Exploration versus exploitation in space, mind, and society [Publisher: NIH Public Access]. Trends in cognitive sciences, 19 (1), 46. 10.1016/j.tics.2014.10.004

Marzilli Ericson, K. M., White, J. M., Laibson, D., & Cohen, J. D. (2015). Money earlier or later? simple heuristics explain intertemporal choices better than delay discounting does [Publisher: SAGE Publications Inc]. Psychological Science, 26 (6), 826–833. 10.1177/0956797615572232

Carter, E. C., & Redish, A. D. (2016). Rats value time differently on equivalent foraging and delaydiscounting tasks [Place: US Publisher: American Psychological Association]. Journal of Experimental Psychology: General, 145 (9), 1093–1101. 10.1037/xge0000196

Culbreth, A., Westbrook, A., & Barch, D. (2016). Negative symptoms are associated with an increased subjective cost of cognitive effort. Journal of abnormal psychology, 125 (4), 528–536. 10.1037/abn0000153

Geana, A., Wilson, R., Daw, N. D., & Cohen, J. (2016). Information-seeking, learning and the marginal value theorem: A normative approach to adaptive exploration. CogSci.

Gillan, C. M., Kosinski, M., Whelan, R., Phelps, E. A., & Daw, N. D. (2016). Characterizing a psychiatric symptom dimension related to deficits in goal-directed control. eLife, 5, e11305. 10.7554/eLife.11305

Ang, Y.-S., Lockwood, P., Apps, M. A. J., Muhammed, K., & Husain, M. (2017). Distinct subtypes of apathy revealed by the apathy motivation index. PloS One, 12 (1), e0169938. 10.1371/journal.pone.0169938

Chong, T., Apps, M., Giehl, K., Sillence, A., Grima, L. L., & Husain, M. (2017). Neurocomputational mechanisms underlying subjective valuation of effort costs. PLoS biology, 15 (2), e1002598. 10.1371/journal.pbio.1002598

Constantino, S. M., Dalrymple, J., Gilbert, R. W., Varanese, S., Di Rocco, A., & Daw, N. D. (2017, August 8). A neural mechanism for the opportunity cost of time (preprint). Neuroscience. 10.1101/173443

Lenow, J. K., Constantino, S. M., Daw, N. D., & Phelps, E. A. (2017). Chronic and acute stress promote overexploitation in serial decision making. The Journal of Neuroscience: The Official Journal of the Society for Neuroscience, 37 (23), 5681–5689. 10.1523/JNEUROSCI.3618-16.2017

Shenhav, A., Musslick, S., Lieder, F., Kool, W., Griffiths, T. L., Cohen, J. D., & Botvinick, M. M. (2017). Toward a rational and mechanistic account of mental effort [eprint: 10.1146/annurev-neuro-072116-031526]. Annual Review of Neuroscience, 40 (1), 99–124. 10.1146/annurev-neuro-072116-031526

Zald, D. H., & Treadway, M. T. (2017). Reward processing, neuroeconomics, and psychopathology. Annual Review of Clinical Psychology, 13, 471–495. 10.1146/annurev-clinpsy-032816-044957

Grahek, I., Everaert, J., Krebs, R. M., & Koster, E. H. W. (2018). Cognitive control in depression: Toward clinical models informed by cognitive neuroscience [Publisher: SAGE Publications Inc]. Clinical Psychological Science, 6 (4), 464–480. 10.1177/2167702618758969

Hayden, B. Y. (2018). Economic choice: The foraging perspective. Current Opinion in Behavioral Sciences, 24, 1–6. 10.1016/j.cobeha.2017.12.002

Inzlicht, M., Shenhav, A., & Olivola, C. Y. (2018). The effort paradox: Effort is both costly and valued. Trends in Cognitive Sciences, 22 (4), 337–349. 10.1016/j.tics.2018.01.007

Juvina, I., Nador, J., Larue, O., Green, R., Harel, A., & Minnery, B. S. (2018). Measuring individual differences in cognitive effort avoidance. In T. Rogers, M. Rau, X. Zhu, & C. Kalish (Eds.), Proceedings of the 40th annual conference of the cognitive science society (pp. 1886–91).

Lopez-Gamundi, P., & Wardle, M. C. (2018). The cognitive effort expenditure for rewards task (c-EEfRT): A novel measure of willingness to expend cognitive effort. Psychological Assessment, 30 (9), 1237–1248. 10.1037/pas0000563

Marchetti, I., Shumake, J., Grahek, I., & Koster, E. H. W. (2018). Temperamental factors in remitted depression: The role of effortful control and attentional mechanisms. Journal of Affective Disorders, 235, 499–505. 10.1016/j.jad.2018.04.064

Musslick, S., Cohen, J. D., & Shenhav, A. (2018). Estimating the costs of cognitive control from task performance: Theoretical validation and potential pitfalls, 6.

Salamone, J. D., Correa, M., Yang, J.-H., Rotolo, R., & Presby, R. (2018). Dopamine, effort-based choice, and behavioral economics: Basic and translational research. Frontiers in Behavioral Neuroscience, 12. Retrieved May 30, 2023, from https://www.frontiersin.org/articles/10.3389/fnbeh.2018.00052

Agrawal, M., Mattar, M., Daw, N., & Cohen, J. D. (2019). Rational arbitration of hippocampal replay. undefined. Retrieved November 1, 2022, from https://www.semanticscholar.org/paper/Rational-Arbitration-of-Hippocampal-Replay-Agrawal-Mattar/b42f4eae895d2c7c6dd634492c7ea9cf433f5e2c

Eisenberg, I. W., Bissett, P. G., Enkavi, A. Z., Li, J., MacKinnon, D. P., Marsch, L. A., & Poldrack, R. A. (2019). Uncovering the structure of self-regulation through data-driven ontology discovery — nature communications. Nature Communications, 10 (1), 1–13. 10.1038/s41467-019-10301-1

Giboin, L.-S., & Wolff, W. (2019). The effect of ego depletion or mental fatigue on subsequent physical endurance performance: A meta-analysis. Performance Enhancement & Health, 7 (1), 100150. 10.1016/j.peh.2019.100150

Grahek, I., Shenhav, A., Musslick, S., Krebs, R. M., & Koster, E. H. W. (2019). Motivation and cognitive control in depression. Neuroscience & Biobehavioral Reviews, 102, 371–381. 10.1016/j.neubiorev.2019.04.011

Patzelt, E. H., Kool, W., Millner, A. J., & Gershman, S. J. (2019). The transdiagnostic structure of mental effort avoidance [Number: 1 Publisher: Nature Publishing Group]. Scientific Reports, 9 (1), 1689. 10.1038/s41598-018-37802-1

Borderies, N., Bornert, P., Gilardeau, S., & Bouret, S. (2020). Pharmacological evidence for the implication of noradrenaline in effort. PLoS Biology, 18 (10), e3000793. 10.1371/journal.pbio.3000793

Garrett, N., & Daw, N. D. (2020). Biased belief updating and suboptimal choice in foraging decisions [Number: 1 Publisher: Nature Publishing Group]. Nature Communications, 11 (1), 3417. 10.1038/s41467-020-16964-5

Strobel, A., Wieder, G., Paulus, P. C., Ott, F., Pannasch, S., Kiebel, S. J., & Kührt, C. (2020). Dispositional cognitive effort investment and behavioral demand avoidance: Are they related? [Publisher: Public Library of Science]. PLOS ONE, 15 (10), e0239817. 10.1371/journal.pone.0239817

Tran, T., Hagen, A. E. F., Hollenstein, T., & Bowie, C. R. (2020). Physical- and cognitive-effort-based decision-making in depression: Relationships to symptoms and functioning: [Publisher: SAGE PublicationsSage CA: Los Angeles, CA]. Clinical Psychological Science. 10.1177/2167702620949236

Wang, H.-T., Smallwood, J., Mourao-Miranda, J., Xia, C. H., Satterthwaite, T. D., Bassett, D. S., & Bzdok, D. (2020). Finding the needle in a high-dimensional haystack: Canonical correlation analysis for neuroscientists. NeuroImage, 216, 116745. 10.1016/j.neuroimage.2020.116745

Bornert, P., & Bouret, S. (2021). Locus coeruleus neurons encode the subjective difficulty of triggering and executing actions [Publisher: Public Library of Science]. PLOS Biology, 19 (12), e3001487. 10.1371/journal.pbio.3001487

Gonález, I., & Déjean, S. (2021, March 1). CCA: Canonical correlation analysis (Version 1.2.1). Retrieved February 23, 2022, from https://CRAN.R-project.org/package=CCA

Hayden, B. Y., & Niv, Y. (2021). The case against economic values in the orbitofrontal cortex (or anywhere else in the brain) [Place: US Publisher: American Psychological Association]. Behavioral Neuroscience, 135 (2), 192–201. 10.1037/bne0000448

Kane, G. A., James, M. H., Shenhav, A., Daw, N. D., Cohen, J. D., & Aston-Jones, G. (2021, September 26). Rat anterior cingulate cortex continuously signals decision variables in a patch foraging task [Section: New Results Type: article]. bioRxiv. 10.1101/2021.06.07.447464

Kilpatrick, Z. P., Davidson, J. D., & El Hady, A. (2021). Uncertainty drives deviations in normative foraging decision strategies. Journal of The Royal Society Interface, 18 (180), 20210337. 10.1098/rsif.2021.0337

Krönke, K.-M., Mohr, H., Wolff, M., Kräplin, A., Smolka, M. N., Bühringer, G., Ruge, H., & Goschke, T. (2021). Real-life self-control is predicted by parietal activity during preference decision making: A brain decoding analysis. Cognitive, Affective, & Behavioral Neuroscience, 21 (5), 936–947. 10.3758/s13415-021-00913-w

Lopez-Gamundi, P., Yao, Y.-W., Chong, T. T.-J., Heekeren, H. R., Mas-Herrero, E., & Marco-Pallaŕes, J. (2021). The neural basis of effort valuation: A meta-analysis of functional magnetic resonance imaging studies. Neuroscience and Biobehavioral Reviews, 131, 1275–1287. 10.1016/j.neubiorev.2021.10.024

Moutoussis, M., Garźon, B., Neufeld, S., Bach, D. R., Rigoli, F., Goodyer, I., Bullmore, E., NSPN Consortium, Guitart-Masip, M., & Dolan, R. J. (2021). Decision-making ability, psychopathology, and brain connectivity. Neuron, 109 (12), 2025–2040.e7. 10.1016/j.neuron.2021.04.019

Musslick, S., & Cohen, J. D. (2021). Rationalizing constraints on the capacity for cognitive control [Publisher: Elsevier]. Trends in Cognitive Sciences, 25 (9), 757–775. 10.1016/j.tics.2021.06.001

Stan. (2021). CmdStanR: The r interface to CmdStan — cmdstanr-package. Retrieved November 2, 2022, from https://mc-stan.org/cmdstanr/reference/cmdstanr-package.html

Toro-Serey, C., Kane, G. A., & McGuire, J. T. (2021). Choices favoring cognitive effort in a foraging environment decrease when multiple forms of effort and delay are interleaved. Cognitive, Affective, & Behavioral Neuroscience. 10.3758/s13415-021-00972-z

Tran, T., Hagen, A. E. F., Hollenstein, T., & Bowie, C. R. (2021). Physical- and cognitive-effort-based decision-making in depression: Relationships to symptoms and functioning [Publisher: SAGE Publications Inc]. Clinical Psychological Science, 9 (1), 53–67. 10.1177/2167702620949236

Zorowitz, S., Niv, Y., & Bennett, D. (2021, April 12). Inattentive responding can induce spurious associations between task behavior and symptom measures. 10.31234/osf.io/rynhk

Agrawal, M., Mattar, M. G., Cohen, J. D., & Daw, N. D. (2022). The temporal dynamics of opportunity costs: A normative account of cognitive fatigue and boredom [Place: US Publisher: American Psychological Association]. Psychological Review, 129, 564–585. 10.1037/rev0000309

Crawford, J. L., Eisenstein, S. A., Peelle, J. E., & Braver, T. S. (2022). Domain-general cognitive motivation: Evidence from economic decision-making – final registered report. Cognitive Research: Principles and Implications, 7 (1), 23. 10.1186/s41235-022-00363-z

Vinckier, F., Jaffre, C., Gauthier, C., Smajda, S., Abdel-Ahad, P., Le Bouc, R., Daunizeau, J., Fefeu, M., Borderies, N., Plaze, M., Gaillard, R., & Pessiglione, M. (2022). Elevated effort cost identified by computational modeling as a distinctive feature explaining multiple behaviors in patients with depression. Biological Psychiatry. Cognitive Neuroscience and Neuroimaging, 7 (11), 1158–1169. 10.1016/j.bpsc.2022.07.011

Westbrook, A., Yang, X., Bylsma, L. M., Daches, S., George, C. J., Seidman, A. J., Jennings, J. R., & Kovacs, M. (2022). Economic choice and heart rate fractal scaling indicate that cognitive effort is reduced by depression and boosted by sad mood. Biological Psychiatry: Cognitive Neuroscience and Neuroimaging. 10.1016/j.bpsc.2022.07.008

Zorowitz, S., & Bennett, D. (2022). NivTurk (Version (v1.2-prolific)). Zenodo. 10.5281/zenodo.6609218

Crawford, J. L., English, T., & Braver, T. S. (2023). Cognitive effort-based decision-making across experimental and daily life indices in younger and older adults. The Journals of Gerontology: Series B, 78 (1), 40–50. 10.1093/geronb/gbac167

Herńan, M. A., & Monge, S. (2023). Selection bias due to conditioning on a collider [Publisher: British Medical Journal Publishing Group Section: Research Methods &amp; Reporting]. BMJ, 381, p1135. 10.1136/bmj.p1135

## References

Mac Donald, A. P. (1970). Revised Scale for Ambiguity Tolerance: Reliability and Validity [Publisher: SAGE Publications Inc]. Psychological Reports, 26 (3), 791–798. 10.2466/pr0.1970.26.3.791

Cacioppo, J. T., & Petty, R. E. (1982). The need for cognition [Place: US Publisher: American Psychological Association]. Journal of Personality and Social Psychology, 42 (1), 116–131. 10.1037/0022-3514.42.1.116

Cacioppo, J. T., Petty, R. E., & Kao, C. F. (1984). The efficient assessment of need for cognition. Journal of Personality Assessment, 48 (3), 306–307. 10.1207/s15327752jpa4803_13

Carver, C. S., & White, T. L. (1994). Behavioral inhibition, behavioral activation, and affective responses to impending reward and punishment: The BIS/BAS Scales [Place: US Publisher: American Psychological Association]. Journal of Personality and Social Psychology, 67 (2),319–333. 10.1037/0022-3514.67.2.319

Cohen, J. D., Forman, S. D., Braver, T. S., Casey, B. J., Servan-Schreiber, D., & Noll, D. C. (1994). Activation of the prefrontal cortex in a nonspatial working memory task with functional MRI. Human Brain Mapping, 1 (4), 293–304. 10.1002/hbm.460010407

Snaith, R. P., Hamilton, M., Morley, S., Humayan, A., Hargreaves, D., & Trigwell, P. (1995). A scale for the assessment of hedonic tone the Snaith-Hamilton Pleasure Scale. The British Journal of Psychiatry: The Journal of Mental Science, 167 (1), 99–103. 10.1192/bjp.167.1.99

Triandis, H. C., & Gelfand, M. J. (1998). Converging measurement of horizontal and vertical individualism and collectivism [Place: US Publisher: American Psychological Association]. Journal of Personality and Social Psychology, 74 (1), 118–128. 10.1037/0022-3514.74.1.118

Kasch, K. L., Rottenberg, J., Arnow, B. A., & Gotlib, I. H. (2002). Behavioral activation and inhibition systems and the severity and course of depression [Place: US Publisher: American Psychological Association]. Journal of Abnormal Psychology, 111 (4), 589–597. 10.1037/0021-843X.111.4.589

McFarland, B. R., Shankman, S. A., Tenke, C. E., Bruder, G. E., & Klein, D. N. (2006). Behavioral activation system deficits predict the six-month course of depression. Journal of Affective Disorders, 91 (2), 229–234. 10.1016/j.jad.2006.01.012

Pinto-Meza, A., Caseras, X., Soler, J., Puigdemont, D., Pérez, V., & Torrubia, R. (2006). Behavioural inhibition and behavioural activation systems in current and recovered major depression participants [Place: Netherlands Publisher: Elsevier Science]. Personality and Individual Differences, 40 (2), 215–226. 10.1016/j.paid.2005.06.021

Cella, D., Yount, S., Rothrock, N., Gershon, R., Cook, K., Reeve, B., Ader, D., Fries, J. F., Bruce, B., & Rose, M. (2007). The Patient-Reported Outcomes Measurement Information System (PROMIS). Medical care, 45 (5 Suppl 1), S3–S11. 10.1097/01.mlr.0000258615.42478.55

Alloy, L. B., Abramson, L. Y., Walshaw, P. D., Cogswell, A., Grandin, L. D., Hughes, M. E., Iacoviello, B. M., Whitehouse, W. G., Urosevic, S., Nusslock, R., & Hogan, M. E. (2008). Behavioral Approach System and Behavioral Inhibition System sensitivities and bipolar spectrum disorders: Prospective prediction of bipolar mood episodes. Bipolar Disorders, 10 (2), 310–322. 10.1111/j.1399-5618.2007.00547.x

Westbrook, A., Kester, D., & Braver, T. S. (2013). What is the subjective cost of cognitive effort? Load, trait, and aging effects revealed by economic preference. PloS One, 8 (7), e68210. 10.1371/journal.pone.0068210

Quilty, L. C., Mackew, L., & Bagby, R. M. (2014). Distinct profiles of behavioral inhibition and activation system sensitivity in unipolar vs. bipolar mood disorders. Psychiatry Research, 219 (1), 228–231. 10.1016/j.psychres.2014.05.007

Costumero, V., Barrós-Loscertales, A., Fuentes, P., Rosell-Negre, P., Bustamante, J. C., & Ávila, C. (2016). BAS-drive trait modulates dorsomedial striatum activity during reward response-outcome associations. Brain Imaging and Behavior, 10 (3), 869–879. 10.1007/s11682-015-9466-5

Pagliaccio, D., Luking, K. R., Anokhin, A. P., Gotlib, I. H., Hayden, E. P., Olino, T. M., Peng, C.-Z., Hajcak, G., & Barch, D. M. (2016). Revising the BIS/BAS Scale to study development: Measurement invariance and normative effects of age and sex from childhood through adulthood [Place: US Publisher: American Psychological Association]. Psychological Assessment, 28 (4), 429–442. 10.1037/pas0000186

Ang, Y.-S., Lockwood, P., Apps, M. A. J., Muhammed, K., & Husain, M. (2017). Distinct Subtypes of Apathy Revealed by the Apathy Motivation Index. PloS One, 12 (1), e0169938. 10.1371/journal.pone.0169938

Constantino, S. M., Dalrymple, J., Gilbert, R. W., Varanese, S., Di Rocco, A., & Daw, N. D. (2017). A Neural Mechanism for the Opportunity Cost of Time (preprint). Neuroscience. 10.1101/173443

Lenow, J. K., Constantino, S. M., Daw, N. D., & Phelps, E. A. (2017). Chronic and Acute Stress Promote Overexploitation in Serial Decision Making. The Journal of Neuroscience: The Official Journal of the Society for Neuroscience, 37 (23), 5681–5689. 10.1523/JNEUROSCI.3618-16.2017

Grahek, I., Everaert, J., Krebs, R. M., & Koster, E. H. W. (2018). Cognitive Control in Depression: Toward Clinical Models Informed by Cognitive Neuroscience [Publisher: SAGE Publications Inc]. Clinical Psychological Science, 6 (4), 464–480. 10.1177/2167702618758969

Le Heron, C., Plant, O., Manohar, S., Ang, Y.-S., Jackson, M., Lennox, G., Hu, M. T., & Husain, M. (2018). Distinct effects of apathy and dopamine on effort-based decision-making in Parkinson’s disease. Brain, 141 (5), 1455–1469. 10.1093/brain/awy110

Kuznetsova, A., Brockhoff, P. B., Christensen, R. H. B., & Jensen, S. P. (2020). lmerTest: Tests in Linear Mixed Effects Models. Retrieved February 25, 2022, from https://CRAN.R-project.org/package=lmerTest

Le Heron, C., Kolling, N., Plant, O., Kienast, A., Janska, R., Ang, Y.-S., Fallon, S., Husain, M., & Apps, M. A. J. (2020). Dopamine modulates dynamic decision-making during foraging [Place: US Publisher: Society for Neuroscience]. The Journal of Neuroscience, 40 (27), 5273–5282. 10.1523/JNEUROSCI.2586-19.2020

Frömer, R., Lin, H., Dean Wolf, C. K., Inzlicht, M., & Shenhav, A. (2021). Expectations of reward and efficacy guide cognitive control allocation [Number: 1 Publisher: Nature Publishing Group]. Nature Communications, 12 (1), 1030. 10.1038/s41467-021-21315-z

Zorowitz, S., Niv, Y., & Bennett, D. (2021). Inattentive responding can induce spurious associations between task behavior and symptom measures. 10.31234/osf.io/rynhk

Bates, D., Maechler, M., Bolker [aut, B., cre, Walker, S., Christensen, R. H. B., Singmann, H., Dai, B., Scheipl, F., Grothendieck, G., Green, P., Fox, J., Bauer, A., & simulate.formula), P. N. K. (c. o. (2022). Lme4: Linear Mixed-Effects Models using ‘Eigen’ and S4. Retrieved March 19, 2022, from https://CRAN.R-project.org/package=lme4

Raio, C. M., Biernacki, K., Kapoor, A., Wengler, K., Bonagura, D., Xue, J., Constantino, S. M., Horga, G., & Konova, A. B. (2022). Suboptimal foraging decisions and involvement of the ventral tegmental area in human opioid addiction [Pages: 2022.03.24.485654 Section: New Results]. 10.1101/2022.03.24.485654

Hernán, M. A., & Monge, S. (2023). Selection bias due to conditioning on a collider [Publisher: British Medical Journal Publishing Group Section: Research Methods &amp; Reporting]. BMJ, 381, p1135. 10.1136/bmj.p1135

